# The Imprinted *Igf2*-*Igf2r* Axis is Critical for Matching Placental Microvasculature Expansion to Fetal Growth

**DOI:** 10.1101/520536

**Authors:** Ionel Sandovici, Aikaterini Georgopoulou, Vicente Pérez-García, Antonia Hufnagel, Jorge López-Tello, Brian Y.H. Lam, Samira N. Schiefer, Chelsea Gaudreau, Fátima Santos, Katharina Hoelle, Giles S.H. Yeo, Keith Burling, Moritz Reiterer, Abigail L. Fowden, Graham J. Burton, Cristina M. Branco, Amanda N. Sferruzzi-Perri, Miguel Constância

**Author notes:** These authors contributed equally (Aikaterini Georgopoulou, Vicente Pérez-García). These authors contributed equally (Cristina M. Branco, Amanda N. Sferruzzi-Perri).

## Abstract

In all eutherian mammals, growth of the fetus is dependent upon a functional placenta, but whether and how the latter adapts to putative fetal signals is currently unknown. Here we demonstrate, through fetal, endothelial, hematopoietic and trophoblast-specific genetic manipulations in the mouse, that endothelial and fetus-derived IGF2 is required for the continuous expansion of the feto-placental microvasculature in late pregnancy. The effects of IGF2 on placental microvasculature expansion are mediated, in part, through IGF2R and angiopoietin-Tie2/TEK signalling. Additionally, IGF2 exerts IGF2R-ERK1/2-dependent pro-proliferative and angiogenic effects on primary feto-placental endothelial cells *ex vivo*. Endothelial and fetus-derived IGF2 also plays an important role in trophoblast morphogenesis, acting through *Gcm1* and *Synb*. Thus, our study reveals a direct role for the imprinted *Igf2-Igf2r* axis on matching placental development to fetal growth and establishes the principle that hormone-like signals from the fetus play important roles in controlling placental microvasculature and trophoblast morphogenesis.

## INTRODUCTION

The mammalian fetus is totally dependent upon the placenta for nutrients and oxygen. Little is known, however, about how placental functional capacity adapts to meet fetal demands for growth. As gestation progresses, the increase in fetal size requires higher levels of nutrients and consequently higher levels of supply *via* placenta. Depending on the species, placental surface area for nutrient exchange increases 5 to 15 fold between mid and late gestation (Fowden et al., 2006). This remarkable adaptation is likely to occur, at least in part, in response to fetus-derived signals, but this important principle remains untested.

We have previously proposed that imprinted genes play central roles in controlling both the fetal demand for, and the placental supply of, maternal nutrients (Constância et al., 2002, Constância et al., 2005, Coan et al., 2008, Angiolini et al., 2011). The *Igf2* (insulin-like growth factor 2) gene encodes a small polypeptide that is highly abundant in both fetal tissues and the fetal circulation. It is one of the most potent growth factors during intrauterine development, affecting the metabolism, proliferation, survival and differentiation of a wide variety of cell types (DeChiara et al., 1991, Baker et al., 1993, Gardner et al., 1999, Burns et al., 2001). In the mouse, homozygous mutants are indistinguishable from growth-deficient littermates with deletion of paternal *Igf2* allele, while mutants with a disrupted maternal *Igf2* allele are phenotypically normal (DeChiara et al., 1991). In humans, reduced *Igf2* expression contributes to the intra-uterine growth restriction in patients with Silver-Russell syndrome (SRS) (Azzi et al., 2014). Conversely, bi-allelic *Igf2* expression caused by loss of *Igf2* imprinting is observed in Beckwith-Wiedemann patients (BWS), a syndrome characterized by somatic overgrowth and increased predisposition to tumours (Azzi et al., 2014).

IGF2 exerts its effects by binding to several IGF/INS receptors (IGF1R, INSR, IGF1/INSR hybrids, IGF2R) (Sferruzzi-Perri et al., 2017). IGF2 binds to IGF2R with the highest affinity, which leads to either IGF2 degradation in the lysosomes or signalling via G-proteins (Okamoto et al., 1990, Maeng et al., 2009, Harris and Westwood, 2012). Additionally, IGF2R has further functions as a mannose 6-phosphate receptor (M6PR) and is also involved in the activation of latent transforming growth factor (TGF)-β1 (Ghosh et al., 2003). In the mouse, *Igf2r* is imprinted, being expressed only from the maternal chromosome (Barlow et al., 1991). Inactivation of the maternal *Igf2r* allele leads to body overgrowth (mutants are ∼30% larger at birth) and perinatal lethality (Lau et al., 1994). This phenotype is largely caused by an excess of extra-cellular IGF2 as shown by the rescue of overgrowth with the introduction of an *Igf2* null allele (Wang et al., 1994). More recently, an IGF2 binding mutant allele (*Igf2r*^I1565A^) was also shown to result in overgrowth and lethality (Hughes et al., 2019). In contrast, imprinting of *IGF2R* in the human is a polymorphic trait, with a minority of cases showing evidence for maternal expression in fetal and/or placental tissues (Xu et al., 1993, Oudejans et al., 2001, Monk et al., 2006).

Here, we apply genetic approaches to define the signalling mechanisms that allow communication between the fetus and placenta, by creating mouse models with a growth mismatch between the two. We first show that circulating IGF2 levels increase in late gestation, with a positive correlation to fetal size. We then provide evidence that endothelial and fetus-derived IGF2 is essential for the appropriate expansion of the feto-placental microvasculature and the underlying trophoblast in late gestation. We also find that endothelial-derived IGF2 plays an essential paracrine role on trophoblast morphogenesis. By contrast, trophoblast-derived IGF2 has only autocrine activities, without any impact on the feto-placental microvasculature. Our work demonstrates that the interaction of circulating IGF2 and endothelial IGF2 with the trophoblast is essential for matching the placental surface area for nutrient exchange to the growth rate of fetal tissues.

## RESULTS

### Expansion of Placental Labyrinthine Zone Coincides with Elevated Levels of Circulating and Endothelial IGF2

The gas and nutrient exchange layer of the mouse placenta (labyrinthine zone – Lz) increased in size with advancing gestational age (Figure 1A), matching the gain in fetal weight (Figure 1B). This is a specific effect of the Lz layer, as placental weight in mice decreases at the very end of gestation (Coan et al., 2004). Concomitantly, fetal plasma IGF2 increased approximately two-fold between E16 and E19 (Figure 1C). At these two developmental stages, we also observed a significant and positive correlation between fetal plasma IGF2 and fetal weights (Figure 1D). Within the placental Lz, *Igf2* expression was highest in feto-placental endothelial cells (FPEC) (Figure 1E) and its mRNA levels increased approximately six-fold between E14 and E19 (Figure 1F). *Igf2* ranked as the highest expressed gene in FPEC RNA-Seq transcriptome at E16, and several other known imprinted genes (Wei et al., 2014) ranked in the top one hundred out of approximately 14,000 genes detected (Figure 1G; Table S1). IGF2 protein was also highly expressed in FPEC (Figure 1H), and significantly higher than in the surrounding trophoblast cells (Figure 1I).

**Figure 1.**
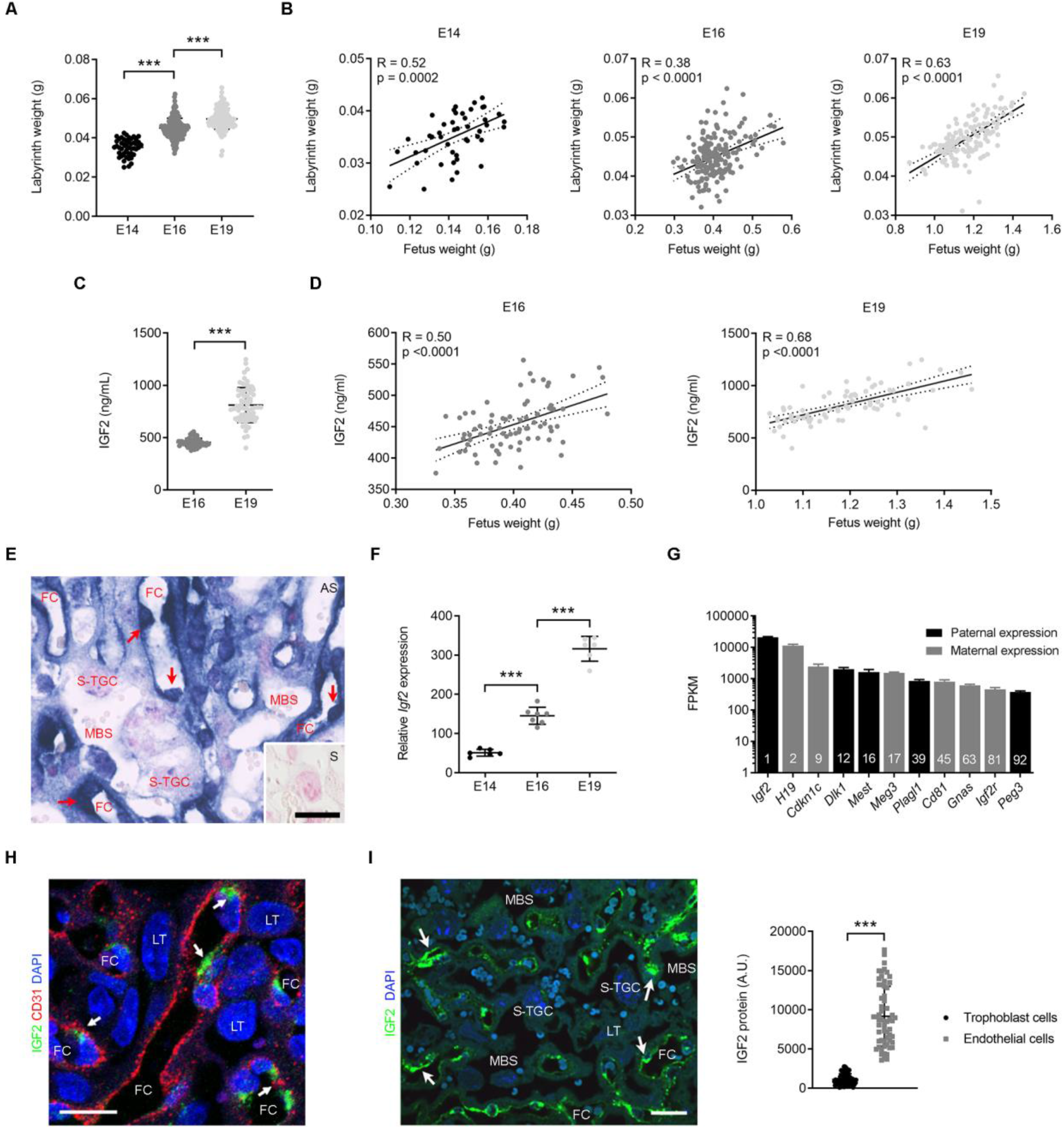
Placental Lz Expansion is Associated with Increasing Levels of Circulating and Endothelial IGF2. (**A**) Weights of micro-dissected labyrinthine zone (Lz). (**B**) Linear correlation analyses between fetal and placental Lz weights [n=46**–**189 placentae from n>10 litters per group in (A) and (B)]. (**C**) Levels of IGF2 (ng/mL) in plasma of wild-type fetuses. (**D**) Linear correlation analyses between fetal weights and circulating IGF2 [n=70**–**79 per group in (C) and (D)]. (**E**) *Igf2* mRNA *in situ* hybridization (blue) in E14 wild-type placental Lz (red arrows – FPEC [feto-placental endothelial cells]; AS – antisense probe; inset with sense probe – S; scale bar is 20µm). (**F**) Relative *Igf2* mRNA expression levels measured by qRT-PCR in FPEC from wild-type placental Lz (n=6–7 per group). (**G**) Imprinted genes that rank within top 100 expressed genes in E16 wild-type FPEC (FPKM – Fragments Per Kilobase Million; n=4). (**H**) Double immunostaining for IGF2 and CD31 in E19 wild-type placenta, demonstrating expression of IGF2 in FPEC. Endothelial cells are very thin and hard to detect except where the cytoplasm is more voluminous around the nucleus, with intense IGF2 stain (white arrows). Transmembrane glycoprotein CD31 immunostaining is in the membrane and largely marks endothelial intercellular junctions (scale bar is 20µm). (**I**) Semi-quantitative measurement of IGF2 protein in FPEC versus trophoblast cells (E19 wild-type placental Lz, n=60 cells per group from two placentae). White arrows – endothelial cells; scale bar is 50µm. For panels (E), (H) and (I) FC – fetal capillaries; MBS – maternal blood spaces; LT – labyrinthine trophoblast cells; S-TGC – sinusoidal trophoblast giant cells. Data in (A), (C), (F), (G) and (I) is presented as averages ± standard deviation (SD); *** *P*<0.001 calculated by one-way ANOVA plus Tukey’s multiple comparisons test in (A) and (F) or by unpaired *t*-test with Welch’s correction in (C) and (I). See also Table S1.

### Fetal and Endothelial IGF2 Control Placental Lz Expansion

To explore whether fetus-derived IGF2 plays a direct role in placental development, we first used a conditional allele (*Igf2*^+/fl^) to delete *Igf2* in the epiblast lineage using the *Meox2*^Cre^ line (Tallquist et al., 2000) (Figures 2A and S1). The deletion of the paternally-inherited *Igf2* allele from embryonic organs and FPEC, but not extra-embryonic tissues, led to placental growth restriction from E14 onwards (Figure 2B). Stereological analyses indicated that only the placental compartments containing embryonic-derived structures (*i.e.* Lz and the chorionic plate – Cp) were smaller in the *Meox2*^Cre/+^ ; *Igf2*^+/fl^ mutants (referred subsequently as *Igf2*^EpiKO^) (Figure 2C). The continuous expansion of the Lz, measured as volume increase between E14 and E19 was severely compromised in mutants (Figure 2C). The overall volume, surface area and total length of fetal capillaries (FC) were normal at E14, but became abnormal from E16 onwards (Figures 2D and S2A). Notably, all other components of the placental Lz, not originating from the embryonic lineage, *i.e.* labyrinthine trophoblast – LT, and maternal blood spaces – MBS, were also reduced in volume to a similar extent as the FC (Figure 2D). These findings provide evidence for a role of fetus-derived IGF2 on the expansion of placental Lz in late gestation.

**Figure 2.**
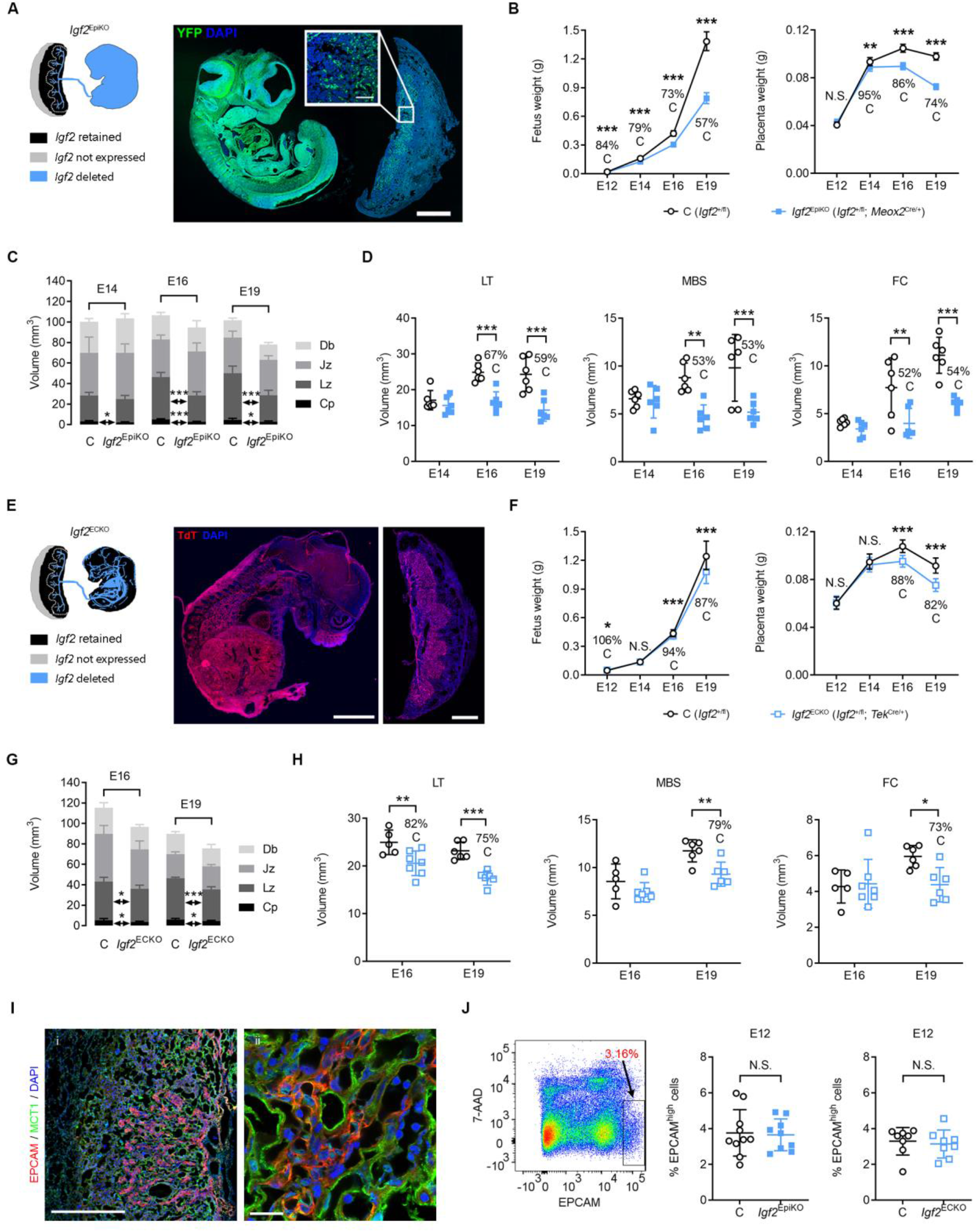
Deletion of *Igf2* in the Epiblast or Endothelium Impairs Placental Lz Expansion. (**A**) Left: schematic of *Igf2* expression in conceptuses with conditional deletion driven by *Meox2*^Cre^. Right: immunostaining for YFP (green) in a representative fetus and placenta paraffin section at E12 of gestation, double transgenic for *Meox2*^Cre^ and *Rosa26* ^fl^STOP^fl^YFP^10^ reporter. YFP expression is observed throughout the fetus, and in the placenta is localized to the Lz and chorionic plate (high magnification inset). Blue – DAPI stain for nuclei; scale bars are 1 mm (low magnification) and 100 µm (high magnification). (**B**) Fetal and placental growth kinetics, measured as average wet-weights for each genotype per litter (E12: n=10 litters [n=41 C and n=32 *Igf2*^EpiKO^]; E14: n=25 litters [n=114 C and n=88 *Igf2*^EpiKO^]; E16: n=37 litters [n=154 C and n=127 *Igf2*^EpiKO^]; E19: n=37 litters [n=164 C and n=121 *Igf2*^EpiKO^]). (**C**) Absolute volumes of the placental layers (Db – decidua basalis, Jz – junctional zone, Lz – labyrinthine zone, Cp – chorionic plate), measured by stereology (n=6 per group). (**D**) Absolute volumes (in mm^3^) of placental Lz components, measured by stereology (LT – labyrinthine trophoblast, MBS – maternal blood spaces, FC – fetal capillaries) (n=6 per group). (**E**) Left: schematic representation of *Igf2* expression in conceptuses with conditional deletion driven by *Tek*^Cre^. Right: representative confocal microscopy of frozen sections from a fetus and its corresponding placenta, double transgenic for *TeK*^Cre^ and Ai9(RCL-tdT) reporter at E16 of gestation. Scale bars are 2 mm (fetus) and 1 mm (placenta). (**F**) Fetal and placental growth kinetics (E12: n=5 litters [n=17 C and n=16 *Igf2*^ECKO^]; E14: n=8 litters [n=26 C and n=34 *Igf2*^ECKO^]; E16: n=13 litters [n=60 C and n=46 *Igf2*^ECKO^]; E19: n=7 litters [n=31 C and n=27 *Igf2*^ECKO^]). (**G**) Absolute volumes of the placental layers measured by stereology (n=5–7 per group). (**H**) Absolute volumes (in mm^3^) of placental Lz components, measured by stereology (n=5–7 per group). (**I**) Double immunostaining for EPCAM (red) and MCT1 (green) in a representative frozen placental section at E12 of gestation. EPCAM expression is observed as clusters of positive cells within the Lz placenta. Blue – DAPI stain for nuclei; scale bars are 500 µm (left panel) and 20 µm (right panel). (**J**) Analysis of EPCAM^high^ positive cells by flow cytometry. Left panel: example of gating used to identify EPCAM^high^ positive cells (the viability dye 7-Aminoactinomycin D [7-AAD] was used to exclude dead cells). Right: quantification of placental EPCAM^high^ positive cells at E12 in conceptuses with conditional *Igf2* deletion driven by *Meox2*^Cre^ (n=10 C and n=9 *Igf2*^EpiKO^ from two litters) or *Tek*^Cre^ (n=8 C and n=8 *Igf2*^ECKO^ from two litters). For all graphs data is shown as averages; error bars represent SD in (C), (D), (G), (H) and (J) or 95% confidence intervals (95%CI) in (B) and (F); N.S. – statistically non-significant; * *P*<0.05; ** *P*<0.01; *** *P*<0.001 calculated by a mixed effects model in (B) and (F) (see Materials and Methods), two-way ANOVA plus Sidak’s multiple comparisons tests in (D) and (H) or unpaired *t*-tests in (C), (G) and (J). See also Figures S1, S2 and S3.

IGF2 is highly expressed in FPEC, as previously shown (Figure 1E–I). Therefore, we next tested whether endothelial-derived IGF2 plays a role in placental development. Paternal *Igf2* allele deletion in the fetal endothelium, including FPEC, using the *Tek*^Cre^ line (Kisanuki et al., 2001) (Figures 2E and S3A–E) led to a moderate, but significant fetal and placental growth restriction, evident from E16 onwards (Figure 2F). Mutant *Tek*^Cre/+^ ; *Igf2*^+/fl^ (referred subsequently as *Igf2*^ECKO^) placentae had reduced volumes of Cp and Lz at both E16 and E19 (Figure 2G), but less striking when compared to *Igf2*^EpiKO^ mutants (Figure 2C). Within the Lz, the LT was reduced at both E16 and E19, while the MBS and FC were comparable to controls at E16, but significantly reduced at E19 (Figures 2H and S2B).

We next tested if IGF2 derived from hematopoietic cells (HC) contributed to the *Igf2*^ECKO^ phenotype, and deleted the paternal *Igf2* allele in the hematopoietic lineage using the *Vav*^iCre^ line (de Boer et al., 2003). Efficient deletion of *Igf2* in HC (referred to subsequently as *Igf2*^HCKO^, Figures S3F and S3G) did not have any significant impact on fetal and placental growth or on Lz expansion between E14 and E19 (Figure S3H). Additionally, *Igf2*^HCKO^ mutant placentae had FC densities similar to that of controls at E19 (Figure S3I).

Although the ‘small’ Lz phenotype is observed in *Igf2*^EpiKO^ and *Igf2*^ECKO^ mutants only in later gestation, it could originate as result of a reduced pool of multipotent labyrinth trophoblast progenitor (LaTP) cells. LaTP cells are detected as clusters of EPCAM^high^ positive cells between E9.5 and E12.5 (Ueno et al., 2013) (Figure 2I). Using flow cytometry analysis at E12, we found no significant difference in the percentage of EPCAM^high^ positive cells between *Igf2*^EpiKO^ and *Igf2*^ECKO^ mutants and their corresponding littermate controls (Figure 2J).

We conclude that the Lz phenotype observed in *Igf2*^EpiKO^ and *Igf2*^ECKO^ mutants is not the consequence of a reduced pool of multipotent LaTP cells due to defective IGF2 signalling in early placental development. However, our data cannot exclude defects in the differentiation potential of the multipotent LaTP cells. The more severe impact on Lz growth observed in *Igf2*^EpiKO^ mutants compared to *Igf2*^ECKO^ mutants also suggests that full placental Lz expansion in late gestation requires both fetus-derived and endothelial-derived IGF2, but not hematopoietic-cell derived IGF2.

### Fetus-derived IGF2 Is Essential for Placental Morphogenesis and Microvasculature Expansion

To uncover molecular signatures associated with the defective placental Lz expansion in *Igf2*^EpiKO^ mutants, we performed an expression microarray analysis in micro-dissected Lz samples at E19, when the Lz expansion and fetal demand for nutrients reach their maximum in absolute terms. Differentially expressed genes (DEG) were enriched in genes implicated in vasculature development and immune responses (Figures 3A, S4A and S4B). We identified a classic molecular signature of impaired angiogenesis – reduced angiopoietin-Tie2/TEK signalling (Augustin et al., 2009) (Figure 3B; Table S2). Lower levels of *Angpt1* and *Tek*, and increased expression of *Angpt2* were validated by qRT-PCR in an independent set of biological samples at E19 and E16, but not at E14. (Figure 3B). Consistent with the well-established roles of the angiopoietin-Tie2/TEK signalling in the control of endothelial cell survival and proliferation (Augustin et al., 2009), placental TUNEL staining revealed a six-fold increase in apoptotic cell frequency in mutants at E16, specifically in the Lz (Figure 3C). To explore directly the identity of the apoptotic cells, we co-stained E16 *Igf2*^EpiKO^ mutant placentae for TUNEL and laminin, a marker for the fetal capillary basement membrane (Milner et al., 1998). Our analysis revealed that the majority of TUNEL+ cells (86.8 ± 4.25%) co-express laminin (Figure 3D), indicating that a large proportion of the apoptotic cells are FPEC. Furthermore, endothelial cell proliferation measured by flow cytometry was significantly reduced at E16 (Figures 3E and S4C), and this finding was confirmed by immunofluorescence (Figure S4D).

**Figure 3.**
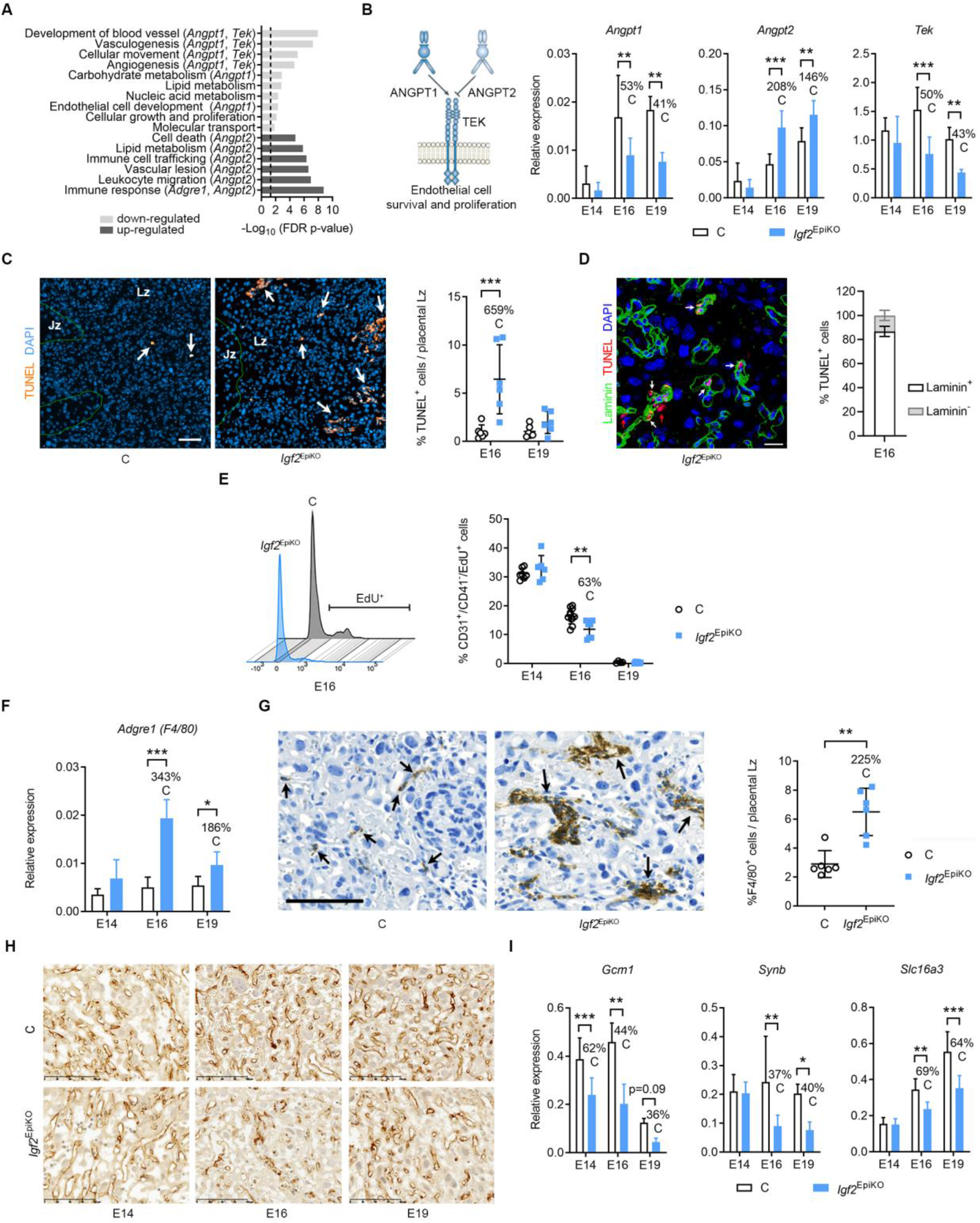
Lack of Fetus-Derived IGF2 Reduces the Expansion of Feto-Placental Microvasculature in Late Gestation. (**A**) Functions enriched in DEGs at E19. (**B**) qRT-PCR analysis of angiopoietin-Tie2/TEK signalling components in placental Lz (n=6–8 per group). (**C**) TUNEL staining in E16 placental Lz (arrows point to apoptotic cells) and data quantification (n=6 samples per group); scale bar is 50µm. (**D**) Left: representative double immunostaining for TUNEL (red) and laminin (green, marker of feto-placental capillaries) in the Lz of an E16 *Igf2*^EpiKO^ mutant placenta (DAPI, blue marks the nuclei; white and red arrows indicate TUNEL^+^ FPECs and LT, respectively; scale bar is 25µm). Right: quantification of TUNEL^+^ cells that are positive or negative for laminin (n=6 *Igf2*^EpiKO^ mutant placentae). (**E**) Feto-placental endothelial cell (FPEC) proliferation measured by flow cytometry (left – representative histograms at E16; right – data quantification; n=4–11 per group). (**F**) qRT-PCR analysis of *Adgre1* in placental Lz. (**G**) Representative F4/80 immunostainings in E16 placental Lz (arrows indicate macrophages). Scale bar is 100µm. Right: percentage of macrophages/placental Lz at E16 (n=6–8 samples per group). (**H**) Representative CD31 immunostaining in placental Lz (scale bar is 100µm). (**I**) qRT-PCR analysis for SynT-II (syncytiotrophoblast layer II) marker genes. For all graphs, data is presented as averages or individual values; error bars are SD; * *P*<0.05, ** *P*<0.01, *** *P*<0.001 by two-way ANOVA plus Sidak’s multiple comparisons tests in (B), (C), (E), (F) and (I) or Mann-Whitney tests in (G). See also Figure S4 and Table S2.

In addition to vascular pathways (which, besides *Angpt1*, *Angpt2* and *Tek*, also include DEGs such as *Dll4*, *Egfl6*, *Fzd4*, *Pdgfc* and *Slit2*, see Table S2), the expression microarrays also identified transcriptional upregulation of genes related to immune responses and leukocyte migration (Figure 3A). Among these was *Adgre1*, a gene that encodes the glycoprotein F4/80, a highly specific cell-surface marker for murine macrophages (Austyn and Gordon, 1981). The up-regulation of *Adgre1* was confirmed by qRT-PCR in placental Lz also at E16 (Figure 3F). Immunostaining for F4/80 showed that the total number of macrophages in Lz was significantly higher in mutants than controls (Figure 3G). Additionally, clusters of macrophages surrounding feto-placental capillaries were found exclusively in mutants (Figure 3G). Next, we assessed the impact of the described increased cell death, reduced cell proliferation and macrophage infiltration, on capillary remodelling across gestation by CD31 immunostaining (marking endothelial cells). The density of FC was dramatically reduced at E16 and E19, suggestive of a disproportionate loss of FPEC (Figure 3H). These CD31-stained or methylene blue-stained resin sections also revealed small areas within the Lz of feto-placental capillaries with accumulations of leucocytes but lacking endothelial cells, or obstructed and thrombotic capillaries surrounded by disorganized and fragmented endothelial cells in late gestation (Figure S4E). Using electron microscopy, we did not observe evidence for feto-maternal barrier interruption that would allow for mixing between maternal and fetal blood, even in areas with disorganized FPECs (Figures S4F and S4G).

Importantly, the array data indicated downregulation of key genes involved in syncytiotrophoblast differentiation (*i.e. Gcm1* and *Synb –* which are expressed specifically in layer II of the syncytiotrophoblast, SynT-II, which is closest to FC; see Table S2). To validate these observations, we performed qRT-PCR and confirmed significant transcriptional reductions across late gestation of SynT-II-specific genes (Rawn and Cross, 2008, Nagai et al., 2010) *Gcm1*, *Synb* and *Slc16a3* (Figure 3I). However, only the SynT-I specific gene *Slc16a1* (Rawn and Cross, 2008, Nagai et al., 2010, Hughes et al., 2013) was modestly down-regulated, but not *Ly6e* and *Syna* (Figure S4H).

Together, our data show that lack of fetus-derived IGF2 triggers dysregulation of angiopoietin-Tie2/TEK signalling in late gestation, with consequent reduced FPEC proliferation and excessive cell death with associated placental macrophage infiltration. It also highlights that fetus-derived IGF2 supports normal development of the trophoblast cells, particularly the SynT-II layer, in a paracrine/endocrine manner, with a knock-on effect on the development of MBS.

### Endocrine IGF2 Is a Fetus-derived Signal that Matches Placental Nutrient Supply Capacity to Fetal Demands for Growth

To provide further insights into the roles of fetus-derived IGF2 in matching placental and fetal growth we analysed five genetic models with either deletion of the paternal *Igf2* allele in fetal tissues, endothelium, trophoblast or ubiquitously, or overexpression of *Igf2* achieved through loss-of-imprinting in fetal tissues (Figure 4). For these models, we used flow cytometry to count FPEC, defined as CD31^+^/CD41^-^ cells (Rhodes et al., 2008 and Figures S5A–C), and measured Lz weight and circulating IGF2 levels. In *Igf2*^EpiKO^ mutants, as expected from the immunostainings shown in Figure 3H, we observed a severe deficit in the total number and the proportion of FPEC at E16 and E19, but normal values at E14 (Figure 4A). The linear Lz expansion expected with gestational age was not observed in this model, matching the severe reductions in FPEC numbers and circulating IGF2 (Figure 4A). In contrast, in *Igf2*^ECKO^ mutants lacking endothelial *Igf2*, circulating levels of IGF2 were only moderately reduced and total numbers of FPEC, but not relative numbers, were only significantly reduced at E19 (Figure 4B). Lz expansion in this model was only blunted at the end of gestation (Figure 4B). A deletion of *Igf2* specifically in the trophoblast cells of the placenta using *Cyp19*^Cre^ (Wenzel and Leone, 2007) (*Igf2*^+/fl^; *Cyp*^Cre/+^ referred subsequently as *Igf2*^TrKO^) (Figures 4C and S6A–E) did not result in changes in FPEC numbers and circulating IGF2, demonstrating that FPEC expansion is independent of trophoblast-derived IGF2. The rate of Lz expansion was normal in this model (Figure 4C). Ubiquitous deletion of *Igf2* in embryo and trophoblast using *CMV*^Cre^ (Schwenk et al., 1995) (*Igf2*^+/fl^; *CMV*^Cre/+^ referred subsequently as *Igf2*^UbKO^) (Figures 4D and S6F) led to a loss of FPEC similar to that observed in the *Igf2*^EpiKO^ mutants, further demonstrating that trophoblast-derived IGF2 does not contribute significantly to FPEC expansion. Lz weight was severely reduced from E14, in line with the near complete absence of IGF2 in fetal circulation (Figure 4D). Conversely, reactivating the transcriptionally silent maternal *Igf2* allele in *H19DMD*^fl/+^ ; *Meox2*^+/Cre^ mutants (Srivastava et al., 2000) (referred subsequently as *H19*-DMD^EpiKO^) (Figures 4E, S6G and S6H), which led to increased levels of circulating IGF2, was associated with an increase of Lz weight and higher numbers of FPEC at E16 and E19 (Figure 4E). Taken together, these results show that IGF2 produced by fetal organs and secreted into the fetal circulation stimulates the expansion of placental Lz, matching FPEC numbers to the fetal demand for growth.

**Figure 4.**
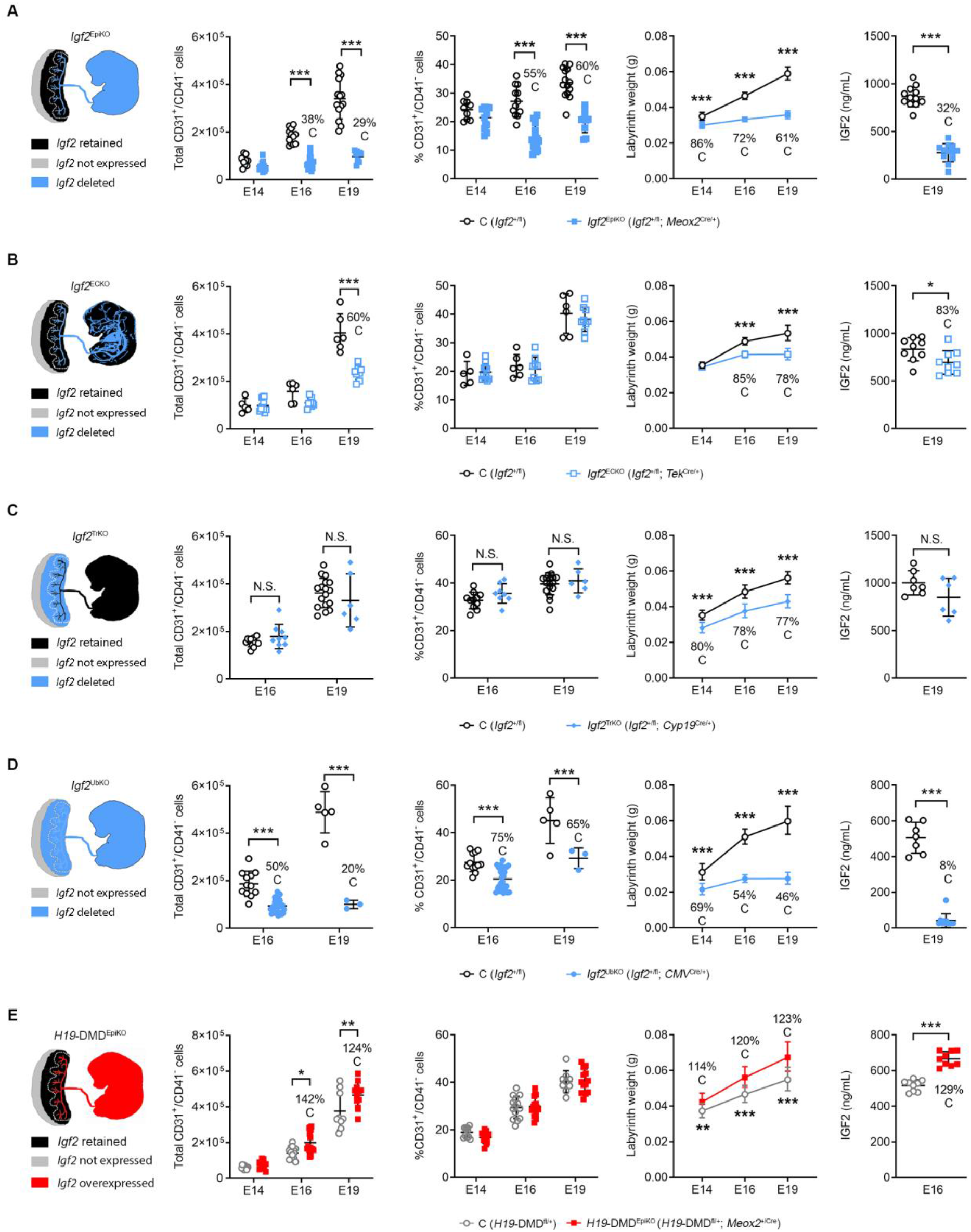
Genetic Models of Mismatched Placental and Fetal Growth Reveal Circulating IGF2 as a Major Endocrine Regulator of FPEC and Placental Lz Expansion. Column 1: schematic diagrams of the genetic models: *Igf2*^EpiKO^ (**A**), *Igf2*^ECKO^ (**B**), *Igf2*^TrKO^ (**C**), *Igf2*^UbKO^ (**D**) *H19*-DMD^EpiKO^ (**E**). Columns 2 and 3: total numbers (column 2) and proportion of FPEC/placental Lz (column 3), measured by flow cytometry (n conceptuses per group: *Igf2*^EpiKO^: n=9–18; *Igf2*^ECKO^: n=5– 11; *Igf2*^TrKO^: n=6–17; *Igf2*^UbKO^: n=3–26; *H19*-DMD^EpiKO^: n=9–15). Column 4: placental Lz growth kinetics (*Igf2*^EpiKO^: n=9–20 litters; *Igf2*^ECKO^: n=3–9 litters; *Igf2*^TrKO^: n=4–9 litters; *Igf2*^UbKO^: n=3–8 litters; *H19*-DMD^EpiKO^: n=3–4 litters). Column 5: IGF2 levels (ng/mL) in plasma (n per group: *Igf2*^EpiKO^: n=12; *Igf2*^ECKO^: n=9; *Igf2*^TrKO^: n=6–7; *Igf2*^UbKO^: n=7–11; *H19*-DMD^EpiKO^: n=9). Data is shown as averages or individual values and error bars are SD (columns 2, 3 and 5) and 95% CI (column 4). N.S. – not significant; * *P*<0.05; ** *P*<0.01; *** *P*<0.001 calculated by two-way ANOVA plus Sidak’s multiple comparisons tests (second and third columns), mixed effects model (fourth column) or Mann Whitney tests (fifth column). See also Figures S5A, S5B and S6.

### IGF2 Signalling Controls Expression of FPEC-derived Angiogenic Factors

To provide further molecular insights into the roles of fetal-derived IGF2 and FEPC-derived IGF2 on microvasculature expansion, we carried out RNA-Seq analysis on FACS-isolated endothelial cells from E16 placental Lz of *Igf2*^EpiKO^ and *Igf2*^ECKO^ mutants and their corresponding controls (Figures 5 and S5C and S5D). Gene ontology (GO) analysis of DEGs identified in *Igf2*^EpiKO^ mutants showed statistical enrichment of biological processes related to immune responses, cell migration, impaired cell proliferation and angiogenesis, extracellular matrix organization and response to hypoxia (Figures 5A and 5B; Table S3). We validated representative DEGs using qRT-PCR in independent biological samples, including genes encoding proteins secreted by the endothelial cells into the extracellular space that have known anti-angiogenic effects [*e.g.*, *Angpt2* (Augustin et al., 2009), *Adamts1* (Lee et al., 2006), *Cxcl10* (Angiolillo et al., 1995) and *Thbs1* (Lawler et al., 2012)], factors implicated in cell migration and response to hypoxia [*Edn1* (Lankhorst et al., 2016)], an interferon-response gene [*Iigp1* (Uthaiah et al., 2003)], an inhibitor of cell proliferation [*Cdkn1a* (Vidal and Koff, 2000)] and a regulator of embryonic vascular development [*Hey2* (Fisher et al., 2004)] (Figure 5C). Importantly, with the notable exception of *Hey2*, all DEGs validated above in the *Igf2*^EpiKO^ model were also identified as DEGs in *Igf2*^ECKO^ mutants, including the up-regulation of *Angpt2*, suggesting that these transcriptional changes are the outcome of autocrine IGF2 actions on FPECs (Figure 5D; Table S3). Next, we searched for transcription factor (TF) binding motifs enriched within the promoters of all DEGs found in the *Igf2*^EpiKO^ model. This analysis identified significant enrichments for binding sites of four TFs encoded by DEGs – KLF4, EGR1, IRF7 and HEY2 (Figure 5E; Table S3). Significantly, the four TFs control the expression of several proteins involved in angiogenesis (labelled with * in Figure 5F and further presented in Table S4), some of which are secreted by the endothelial cells into the extracellular space (Table S4). This analysis also highlighted several chemokines that were up-regulated in FPEC [such as CCL2 (Gregory et al., 2006) and IL15 (Fehniger and Caligiuri, 2001)] that are likely involved in attracting and modulating the activity of macrophages that surround the feto-placental capillaries (as shown in Figure 3G). Thus, based on our data, we propose that IGF2 signalling is necessary for proliferation and survival of FPECs.

**Figure 5.**
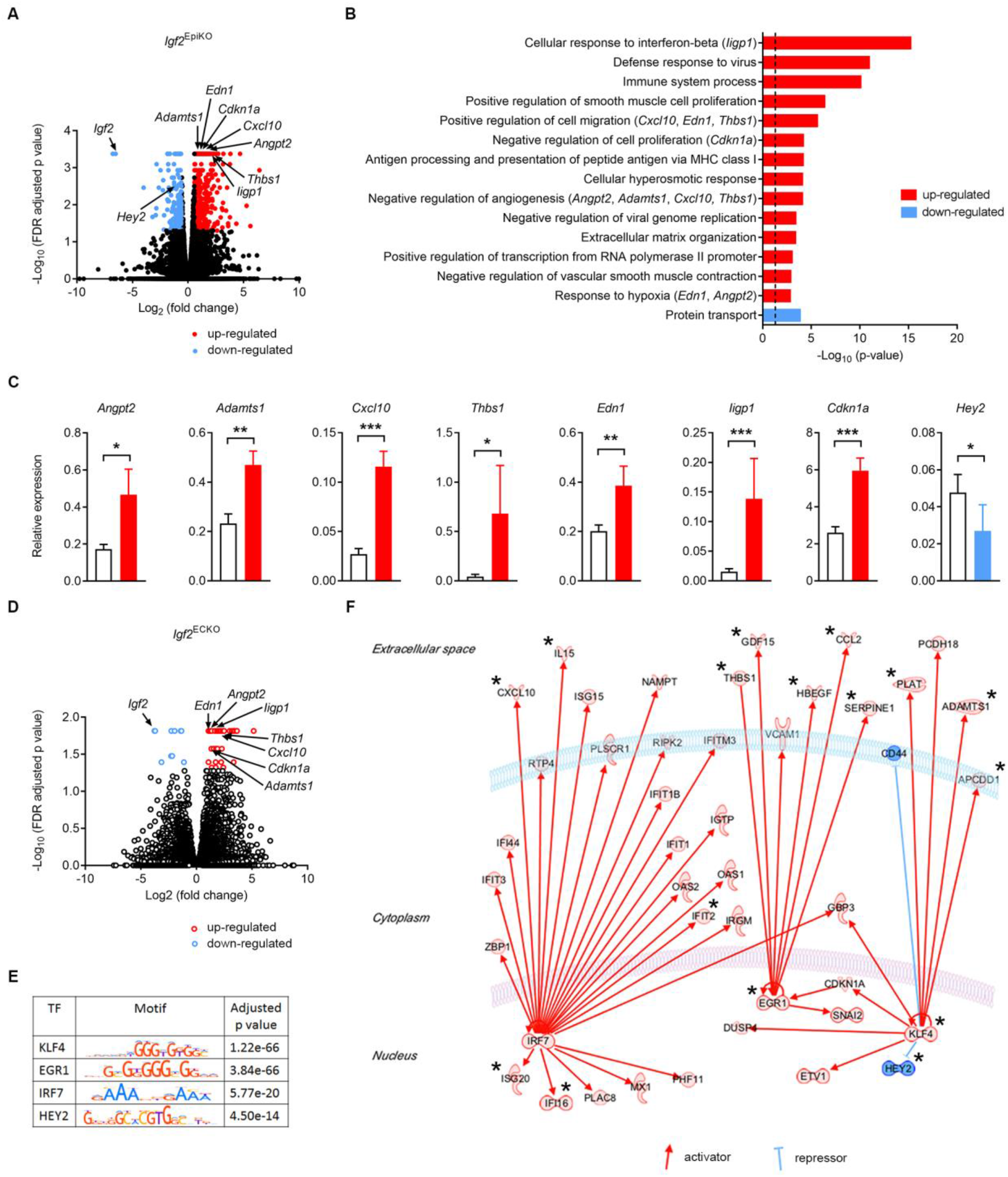
IGF2 Signalling Regulates Angiogenic Properties of Endothelial Cells. (**A**) Volcano plot representation of differentially expressed genes (DEGs) identified by RNA-seq in E16 FPEC (*Igf2*^EpiKO^ versus controls). Significant up-regulated and down-regulated DEGs (FDR<0.05) are shown with red and blue, respectively. (**B**) Top scoring biological processes enriched in DEGs. Biologically validated DEGS are listed in parentheses. The dotted line corresponds to FDR-corrected *P* value of 0.05. (**C**) Biological validation. Data is shown as averages (n=11-12 samples per group); error bars are SEM; * *P*<0.05, ** *P*<0.01, *** *P*<0.001 calculated by Mann-Whitney tests. (**D**) Volcano plot representation of DEGs identified by RNA-seq in E16 FPEC (*Igf2*^ECKO^ versus controls). Significant up-regulated and down-regulated DEGs (FDR<0.05) are shown with red and blue, respectively. (**E**) Transcription factors (TFs) identified by Analysis of Motif Enrichment (AME). (**F**) IPA regulatory network built with the four TFs identified using AME analysis. Proteins labelled with * are known regulators of angiogenesis (angiostatic or pro-angiogenic factors) and key references are listed in Table S4. See also Figures S5C and S5D and Table S3.

### IGF2 Signalling on FPEC Is Mediated by IGF2R *in vitro* and *in vivo*

To further investigate the role of IGF2 in fetal capillary remodelling and identify the receptors that might mediate its effects on endothelial cells, we isolated primary FPEC from E16 wild-type placental Lz and cultured them *ex vivo* (Figures 6A, 6B and 6C). Only the type I (*Igf1r*) and type II (*Igf2r*) receptors were expressed in FPEC both *in vivo* (Figure 6D) and *ex vivo* (Figure 6E). When cultured *ex vivo* for ten days (passage one) FPEC switch off *Igf2* transcription, which differs from FPEC freshly isolated by FACS, (Figure 6F). Exposure of cultured FPEC to exogenous IGF2 significantly increased their ability to form capillary-like tube structures when seeded on matrigel (Figures 6G and 6H), demonstrating that IGF2 exerts direct angiogenic effects on FPEC. We also exposed cultured FPEC to IGF2^Leu27^, an analogue previously shown to bind to IGF2R with high selectivity (Beukers et al., 1991), which stimulated capillary-like tube formation although to a lesser extent compared to IGF2 (Figures 6G and 6H). When FPEC were treated with IGF2 and picropodophyllin (PPP), a small molecule that inhibits phosphorylation of IGF1R without interfering with INSR activity (Girnita et al., 2004), their ability to form capillary-like tube structures was very similar to that of cells treated with IGF2 alone (Figures 6G and 6H). Thus, IGF2 exerts direct angiogenic effects on primary FPEC, which are mediated by IGF2R and are independent of IGF1R.

**Figure 6.**
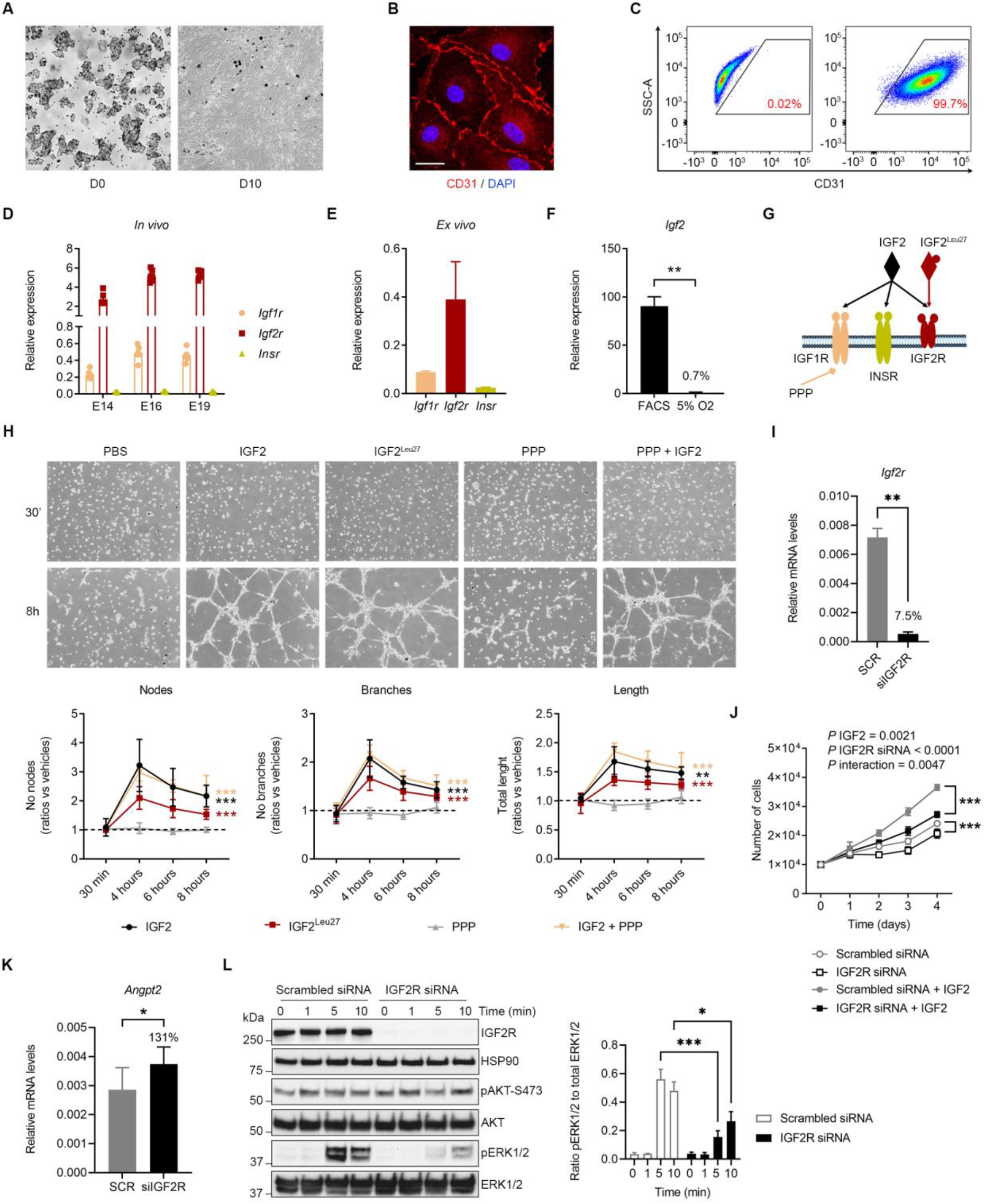
IGF2 Acts on Feto-Placental Endothelial Cells *via* IGF2R-ERK signalling *ex vivo*. (**A**) Primary feto-placental endothelial cell (FPEC) isolated from E16 placental Lz: D0 – freshly isolated cells, bound to magnetic beads coated with anti-CD31 antibodies; D10 – FPEC at passage one (after approximately 10 days of culture). (**B**) Confocal imaging of passage one FPEC, stained for CD31 (scale bar is 20 µm). (**C**) Flow cytometry analysis of passage one FPEC stained for CD31, demonstrating that these are almost exclusively CD31^+^. (**D**) qRT-PCR analysis for *Igf1r*, *Igf2r* and *Insr* in FPECs isolated by FACS (n=6–7 per group). (**E**) Relative expression of the three IGF receptors in passage one FPEC. (**F**) qRT-PCR analysis of *Igf2* mRNA levels in passage one FPEC cultured in 5% O2 versus primary FPEC isolated from E16 placental Lz by FACS. (**G**) Schematic representation of IGF2 and IGF receptors. IGF2^Leu27^ analogue acts specifically on IGF2R and picropodophyllin (PPP) inhibits phosphorylation of IGF1R. (**H**) Representative images of capillary-like tube formation assay in primary FPEC seeded on matrigel and exposed to exogenous IGF2, IGF2^Leu27^, PPP or PPP+IGF2 (equal seeding of cell numbers at 30 min and tube formation at 8 hours), and quantification of number of nodes, branches and total length (n=5–6 independent experiments). (**I**) qRT-PCR analysis of *Igf2r* mRNA levels in primary FPECs upon knockdown by siRNA (n=8 samples/group). (**J**) Proliferation assay of primary FPEC with or without IGF2R siRNA knockdown, in presence or absence of IGF2, on 4 consecutive days after plating. Cells with IGF2R siRNA knockdown exhibit significant proliferation defects that are further accentuated upon IGF2 treatment (n=5 biological replicates per group). (**K**) qRT-PCR analysis of *Angpt2* mRNA levels in primary FPECs transfected with scrambled siRNA or IGF2R siRNA, upon 4 days of treatment with 50 ng/mL mouse recombinant IGF2 (n=8 samples/group). (**L**) Left side: identification of delayed ERK1/2 phosphorylation in FPECs with IGF2R siRNA knockdown upon acute treatment with 50ng/mL mouse recombinant IGF2. HSP90 was used as internal control for protein loading. Right side: quantification of ratios pERK1/2 to total ERK1/2 for n=3 independent biological replicates. For all graphs, data is presented as averages or individual values and error bars represent SEM. * *P*<0.05, ** *P*<0.01 and *** *P*<0.001 calculated by a Mann Whitney test in (F), two-way ANOVA tests with Sidak’s multiple comparisons test in (H), (J) and (L), Wilcoxon matched-pairs signed rank test in (I) and paired student *t*-test in (K).

To investigate further the observed effects of IGF2R in mediating IGF2 actions on primary FPECs, we performed *Igf2r* knockdown using siRNA and investigated the impact on cell proliferation and intracellular signalling with or without exogenous IGF2 stimulation (Figure 6I–L). Upon IGF2 stimulation, efficient *Igf2r* knockdown led to reduced FPEC proliferation, demonstrating that the pro-proliferative actions of IGF2 on FPEC require IGF2R (Figure 6J). However, *Igf2r* knockdown also resulted in reduced FPEC proliferation, even in the absence of IGF2 stimulation, suggesting that IGF2R is required for normal FPEC proliferation, independent of IGF2 (Figure 6J). Additionally, FPEC stimulated with IGF2 for 96 hours, but lacking IGF2R showed significant up-regulation of *Angpt2* mRNA levels (Figure 6K). Acute IGF2 stimulation did not activate AKT, a key signalling node downstream of IGF1R (Figure 6L). AKT phosphorylation was not affected by the IGF2R knockdown (Figure 6K). However, we observed a significant delay in pERK1/2 phosphorylation upon acute stimulation of FPEC with IGF2 (Figure 6K). These data demonstrate both IGF2-independent and IGF2-dependent actions of IGF2R in controlling FPEC proliferation and highlight the role of ERK pathway in mediating the actions of IGF2 on FPEC *via* IGF2R.

We further confirmed these *in vitro* findings using conditional deletions of IGF1R and IGF2R receptors *in vivo*. Accordingly, efficient homozygous deletion of *Igf1r* from the endothelium (*Igf1r*^ECKO^) did not have any significant impact on fetal, whole placenta or placental Lz growth kinetics, nor did it alter the total and relative numbers of FPEC/Lz, apart from a slight increase in the percentage of FPEC at E19 (Figures S7). Strikingly, the deletion of maternally-expressed *Igf2r* allele from the endothelium (*Igf2r*^ECKO^ – see Figures 7A, 7B and 7C) resulted in a reduction in the percentage of FPEC/placental Lz at both E16 and E19 (Figure 7D), further confirmed by a reduced density of CD31^+^ cells by immunofluorescent staining (Figure 7E). The total number of FPEC/Lz was also significantly reduced at E16, but became normal at E19 (Figure 7D). Notably, the overall placenta and Lz were overgrown, from E16 onwards (Figures 7C and 7F), coincident with an increase in levels of circulating IGF2 in plasma (Figure 7G). Together, our *in vitro* and *in vivo* experiments demonstrate that IGF2R mediates, at least partially, the signalling actions of IGF2 on FPEC.

**Figure 7.**
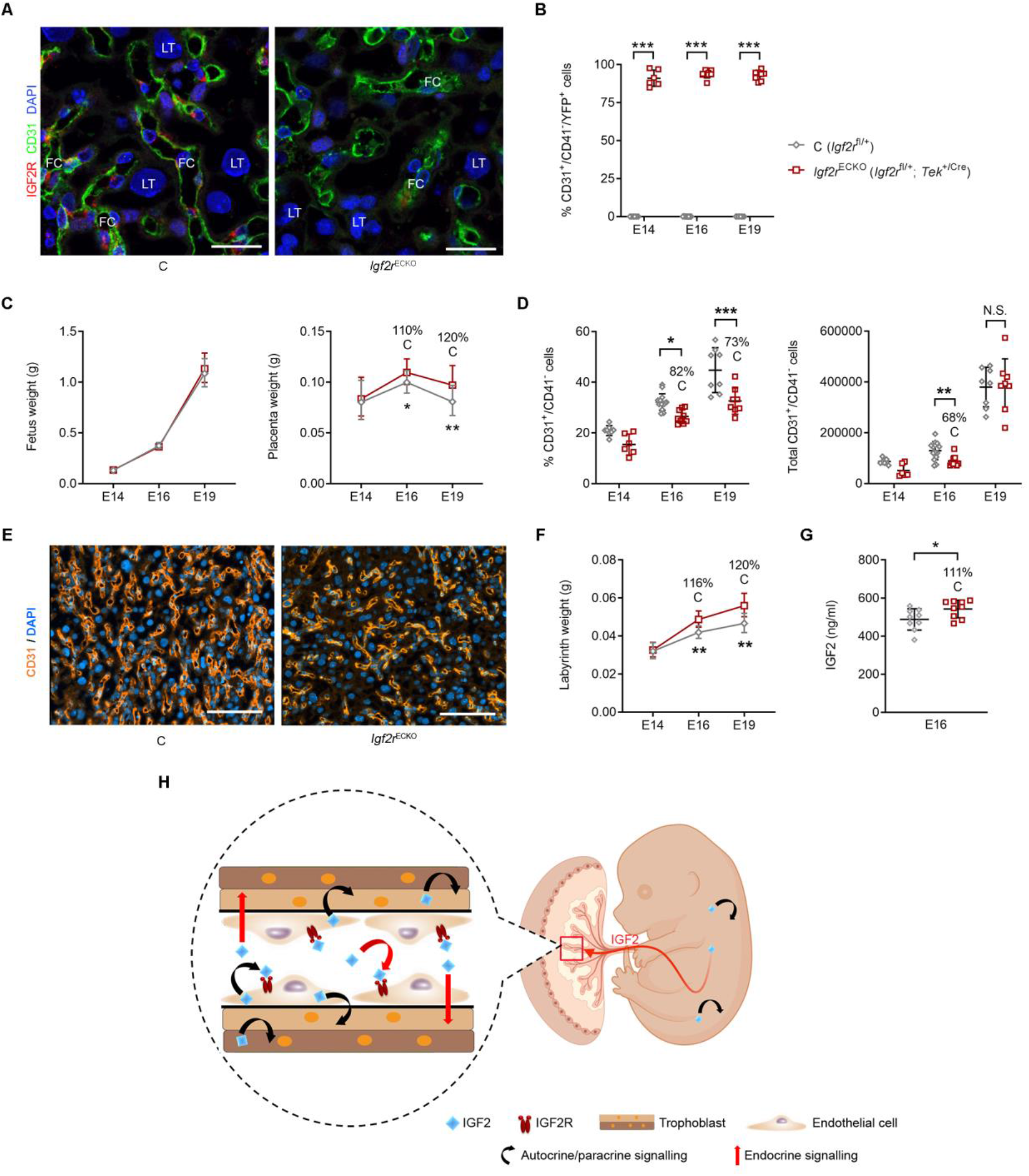
IGF2 Acts on Feto-Placental Endothelial Cells via IGF2R *in vivo*. (**A**) Representative double immunostaining for IGF2R (red) and the endothelial cell marker CD31 (green) in *Igf2r*^ECKO^ mutant and control placental Lz at E16 (DAPI, blue marks the nuclei; FC – fetal capillaries, LT – labyrinthine trophoblast; scale bar is 25µm). (**B**) Flow cytometry analysis showing that the majority (>80%) of *Igf2r*^ECKO^ mutant feto-placental endothelial cells (FPECs) express YFP, demonstrating good efficiency of *Tek2*^Cre^ in these samples (n=6–14 per genotype). (**C**) Fetal and placental growth kinetics in *Igf2r*^ECKO^ (*Igf2r*^fl/+^; *Tek*^+/Cre^) mutants compared to *Igf2r*^fl/+^ controls (n=8–28 conceptuses from n=3-8 litters for each developmental stage). (**D**) Proportion and total numbers of FPEC/placental Lz measured by flow cytometry (n=6–14 per group). (**E**) Representative CD31 immunofluorescence staining in E16 placental Lz (scale bar is 100µm). (**F**) Placental Lz growth kinetics: *Igf2r*^ECKO^ (n=8–16 conceptuses per group). (**G**) IGF2 levels (ng/mL) in plasma at E16 (n=9 per group). (**H**) Model summarizing the proposed actions of fetus-, endothelial- and trophoblast-derived IGF2. For all graphs, data is presented as averages or individual values and error bars represent SD in (B), (D), and (G), or 95%CI in (C) and (F). N.S. – not significant; * *P*<0.05; ** *P*<0.01; *** *P*<0.001 calculated by two-way ANOVA tests in (B), and (D), mixed effects model in (C) and (F) and Mann Whitney tests in (G). See also Figure S7.

## DISCUSSION

The major findings of this study are the identification of the imprinted *Igf2*-*Igf2r* axis as a key pathway that controls the expansion of the placental vascular tree in late gestation, and the demonstration that fetus-derived signals are important regulators of placental development and function. Although a vast number of genetic pathways have been discovered that are important for the development of different cell types in the placenta and the fetus, there are no functional genetic investigations to date on how the fetus signals its nutrient requirements to the placenta and how the placenta matches these demands (for example through the increase in surface area in late gestation). We tackled these questions with an experimental design based on the manipulation of the growth rate of fetal tissues independent of the placenta, and vice-versa, in the mouse. We used conditional targeting of imprinted genes with well-established growth functions (*Igf2*, *Igf2r*, *H19*) as model systems (importantly, due to imprinting, the mother is phenotypically normal). The analysis of these models of mismatch between fetal and placental growth allowed us to establish a number of novel mechanistic principles that regulate the cooperative signalling between the fetus and the placenta and, consequently, the control of maternal resources.

Firstly, we found that circulating IGF2 correlates positively with fetal size in late gestation, reflecting the growth rate of fetal tissues and the demand for nutrients. Mice with a severe decrease in levels of circulating/fetal IGF2 showed a drastic (and disproportionate) loss of feto-placental endothelial cells. This severe placental angiogenesis phenotype was associated with reduced endothelial cell proliferation and increased apoptosis, altered differentiation of the underlying trophoblast and reduced density of MBS, ultimately leading to a failure in the expansion of the Lz and surface area for nutrient transport. Conversely, increased requirements for nutrients caused by bi-allelic *Igf2* expression, which drove higher growth rates, led to ‘overexpansion’ of the Lz. Therefore, we show that greater fetal demands for growth, driven by IGF2, signals enhanced placental growth. Secondly, we also found that FPECs are a significant source of IGF2, with levels increasing with gestational age. Endothelial *Igf2*-deficient mice show ∼17% reduction in circulating IGF2 and impaired expansion of the microvasculature and Lz, but no disproportionate reduction in number of FPECs (which is only seen when circulating IGF2 is severely reduced). These findings strongly suggest that hormone-like signals from the fetus, such as IGF2, are also required for the normal expansion of the Lz and surface area of the placenta. Importantly, we ruled out the hypothesis that failure in expansion is due to a reduction in the number of LaTP. IGF2 has been reported to be an essential component of maintenance of stem-cell niches in other organs (Ferron et al., 2015, Ziegler et al., 2019). Our data suggest that endothelial IGF2 and circulating IGF2 are not required for the proliferation and maintenance of LaTP pools, but rather their differentiation.

Based on the experimental evidence provided in this study, we propose a model (Figure 7H) in which fetus-derived IGF2, from multiple tissues, is the signal that allows matching placental supply capacity to the fetal demands for growth. At the placental interface, circulating IGF2 directly stimulates endothelial cell proliferation and survival, and capillary branching in part through IGF2R (as shown *in vivo* and *ex vivo*). Circulating IGF2 may also directly control the growth and differentiation of the underlying trophoblast, as it can cross (in free form or in binary complexes) the capillary walls or permeate through the fenestrated endothelium (Bach, 2015). We suggest that the feto-placental endothelium is a large reservoir of IGF2, boosting further IGF2 signalling, and acting in a paracrine and autocrine manner to control the growth and remodelling of fetal capillaries, with a ‘secondary effect’ on trophoblast morphogenesis. On the contrary, hematopoietic cells that originate from precursors common to FPECs (Rhodes et al., 2008) do not play any significant role in placental Lz expansion. Importantly, the effect of IGF2 signalling on feto-placental microvascular remodelling seems specifically driven by fetus-derived IGF2. Accordingly, we did not find any evidence that IGF2 produced by the trophoblast has a direct role on vascularization, being instead required for trophoblast morphogenesis. We therefore suggest that the key role of circulating IGF2 is to provide fetus-derived angiogenic signals to promote the vascular tree expansion in later gestation, in conjunction with local IGF2, derived from the fetal endothelium of the placenta. Mechanistically, likely molecular triggers of fetus-derived IGF2 signalling on microvasculature expansion and trophoblast morphogenesis are IGF2R-ERK1/2-angiopoietin-Tie2/TEK signalling and the key trophoblast differentiation genes *Gcm1* and *Synb*, respectively. Activation of ERK1/2 signalling pathway by IGF2 *via* IGF2R has been observed *in vitro* in previous studies (El-Shewy et al., 2006; El-Shewy et al., 2007). Additionally, primary aortic endothelial cells isolated from *Erk1*/*Erk2* double knockout mice associated transcriptional up-regulation of several DEGs identified in the *Igf2*^EpiKO^ FPECs, including *Angpt2*, *Adamts1*, *Igtp* and *Ifit1* (Srinivasan et al., 2009) (see Figure 5). Although the detailed mechanisms by which ERK1/2 signalling leads to changes in *Angpt2* expression remain to be elucidated, these observations are compatible with a model in which IGF2 binds to IGF2R to activate ERK1/2 signalling pathway, leading to lower *Angpt2* expression, as well as other angiostatic factors (described in Figure 5). We found no evidence for de-regulation of known controllers of placenta angiogenesis, such as VEGF (vascular growth endothelial factor) and PGF (placental growth factor) (Aplin et al., 2020) in these mouse models. Instead, the late gestation angiogenesis defects are related to angiopoietins, although the contribution of other pathways cannot be ruled out. Importantly, angiopoietin-Tie2/TEK signalling has also been implicated in trophoblast morphogenesis, independent of their vascular actions (Kappou et al., 2015), and therefore may be an important link between vascular effects and trophoblast in these models. To our knowledge this is the first report of the impact of the vasculature on trophoblast morphogenesis acting in late gestation.

The genetic models used in our study have several limitations. One is the inability to specifically target circulating levels of IGF2 without impacting on the size of the fetus, but also the lack of specific Cre-lines to the placental endothelium. The source of circulating IGF2 is likely to be multi-organ. Unlike for IGF1, the liver is not the main source of circulation IGF2 in mice (Sandovici et al., unpublished). Moreover, and consistent with high expression of IGF2 in mesoderm-derived tissues, deletion of mesodermal *Igf2* enhancers results in reductions in circulating IGF2 by ∼50% (Davies et al., 2002). Future transgenic studies targeting specifically circulating IGF2 or placental endothelium are therefore likely to be very challenging. The overexpression IGF2 models (*H19*-DMD^EpiKO^ and *Igf2r*^ECKO^) used in this study are not without their limitations. Small contributions to the phenotypes due to the actions of the *H19*-encoded *miR-675* and the mannose-6-phosphate receptor roles of IGF2R/activation of TGF-β1 (thus independent of IGF2) cannot be completely ruled out. Indeed, in our study we observed IGF2-independent effects of IGF2R *ex vivo* (reduced basal levels of FPEC proliferation upon siRNA knockdown of *Igf2r*). However, our conclusions that IGF2R is an important IGF2-receptor driving the endothelial cell phenotypes are based on both *in vitro* and *in vivo* evidence. Moreover, *Insr*, and therefore INSR-IGF1R hybrids, are expressed at very low levels in endothelial cells and unlikely to be functional. Endothelial specific deletion of *Igf1r* is not associated with a phenotype. It is important to note that a major target of the *H19*-encoded *mir-675* is IGF1R (Keniry et al., 2012), which is therefore unlikely to be of functional relevance in this context. Importantly, partial redundancy between actions of IGF1R and IGF2R in FPECs cannot be completely ruled out based on this study and warrants future experiments involving dual conditional deletions of *Igf1r* and *Igf2r* driven by *Tek*-Cre.

Our study has a number of important implications. It provides insights into the complex interplay between trophoblast branching morphogenesis and placental vascularization. To our knowledge, IGF2 is the first example of a fetus-derived hormone-like molecule that signals to the placenta and adapts the expansion of feto-placental microvasculature and trophoblast morphogenesis to the embryo size. Matching placental supply capacity to fetal demand for growth also involves IGF2R – the other imprinted member of the IGF family (Constância et al., 2004). The imprinting of the IGF system is thus likely to have played a key evolutionary role in the origins of the expansion of the feto-placental microvasculature and surface area for nutrient transport throughout pregnancy – a fundamental biological process that is observed in all eutherian species (Fowden et al., 2006). In humans, circulating levels of IGF2 in the umbilical cord progressively increase between 29 weeks of gestation and term, similarly to our findings in the mouse (Gohlke et al., 2004). Additionally, large-for-gestational age and small-for-gestational age babies, have been reported to show increased and reduced levels of IGF2 in the umbilical cord, respectively (Verhaeghe et al., 1993, Tzschoppe et al., 2015). Moreover, placentae obtained from imprinting growth syndrome patients with disrupted IGF2 signalling are often associated with placentomegaly in BWS cases, due to hypervascularization and hyperplasia (Aoki et al., 2011, Armes et al., 2012) and small hypoplastic placentas in SRS cases (Yamazawa et al., 2008), showing striking similarities to our mouse studies. Importantly, most cases of poor placentation in FGR (fetal growth restriction) reported so far were related to placental malperfusion from the maternal side and in response to a perturbed maternal environment (Mayhew et al., 2004). Our findings suggest that poor placentation in humans could be caused by deficient microvasculature expansion due to reduced fetus-derived IGF2 signalling, with important clinical implications.

## ACKNOWLEDGEMENTS

We thank Matt Castle (GSLS Biostatistics Initiative, University of Cambridge) and Wendy Cooper for help with statistical analyses, Jeremy Skepper, Nuala Daw, Barbara Villela and Bliss Anderson for technical assistance with placental stereology and transmission electron microscopy analyses, Adrian Wayman, Laura Hunter (West Forvie Phenomics Center) and Edina Gulacsi (Sferruzzi-Perri laboratory) for help with mouse husbandry; Keli Philips and James Warner (Histology Core Facility) for help with preparing tissue samples for histology and F4/80 staining, Gregory Strachan (Imaging Core Facility) for help with TUNEL^+^ and F4/80^+^ cells counting using HALO, Marcella Ma (Genomics and Transcriptomics Core) for help with preparing the RNA-Seq libraries; Evgeniya Shmeleva and Francesco Colucci for advice regarding flow cytometry analyses and providing aliquots of several antibodies used for flow cytometry and FACS; Natalia Savinykh and Esther Perez (NIHR Cambridge BRC Cell Phenotyping Hub) for help with flow cytometry cell sorting. **Funding:** This work was supported by Biotechnology and Biological Sciences Research Council (grant BB/H003312/1 to M.C.), Medical Research Council (MRC_MC_UU_12012/4 to M.C.; MRC_MC_UU_12012/5 to the MRC Metabolic Diseases Unit; MR/R022690/1 to A.N.S-P.), Spanish Ministry of Science and Innovation (RYC-2019-026956 and PID2020-114459RA-I00 to V.P-G.), Royal Society (Dorothy Hodgkin Research Fellowship grant DH130036 to A.N.S-P.), Centre for Trophoblast Research and the NIHR Cambridge BRC Cell Phenotyping Hub.

## AUTHOR CONTRIBUTIONS

I.S. and A.G. performed all the *in vivo* experimental work, with contributions from A.H., J.L-T., S.N.S., F.S., K.H. and A.N.S-P. I.S., B.Y.H.L. and G.S.H.Y. performed bioinformatics analyses. I.S., M.R. and C.M.B. performed the *in vitro* tube formation assays. V.P-G. performed the *in vitro Igf2r* knockdown experiments on primary placental endothelial cells. K.B. developed and performed the assay for IGF2 measurements in fetal plasma. I.S. and M.C. designed the project and G.J.B., A.L.F., A.N.S-P. and C.M.B. assisted with the experimental design and data analysis/interpretation. I.S., G.J.B. and M.C. wrote the manuscript, with important contributions from A.L.F., C.M.B. and A.N.S-P. All other authors discussed the results and edited the manuscript. M.C. managed and supervised all aspects of the study.

## STAR*METHODS

### Key Resources Table

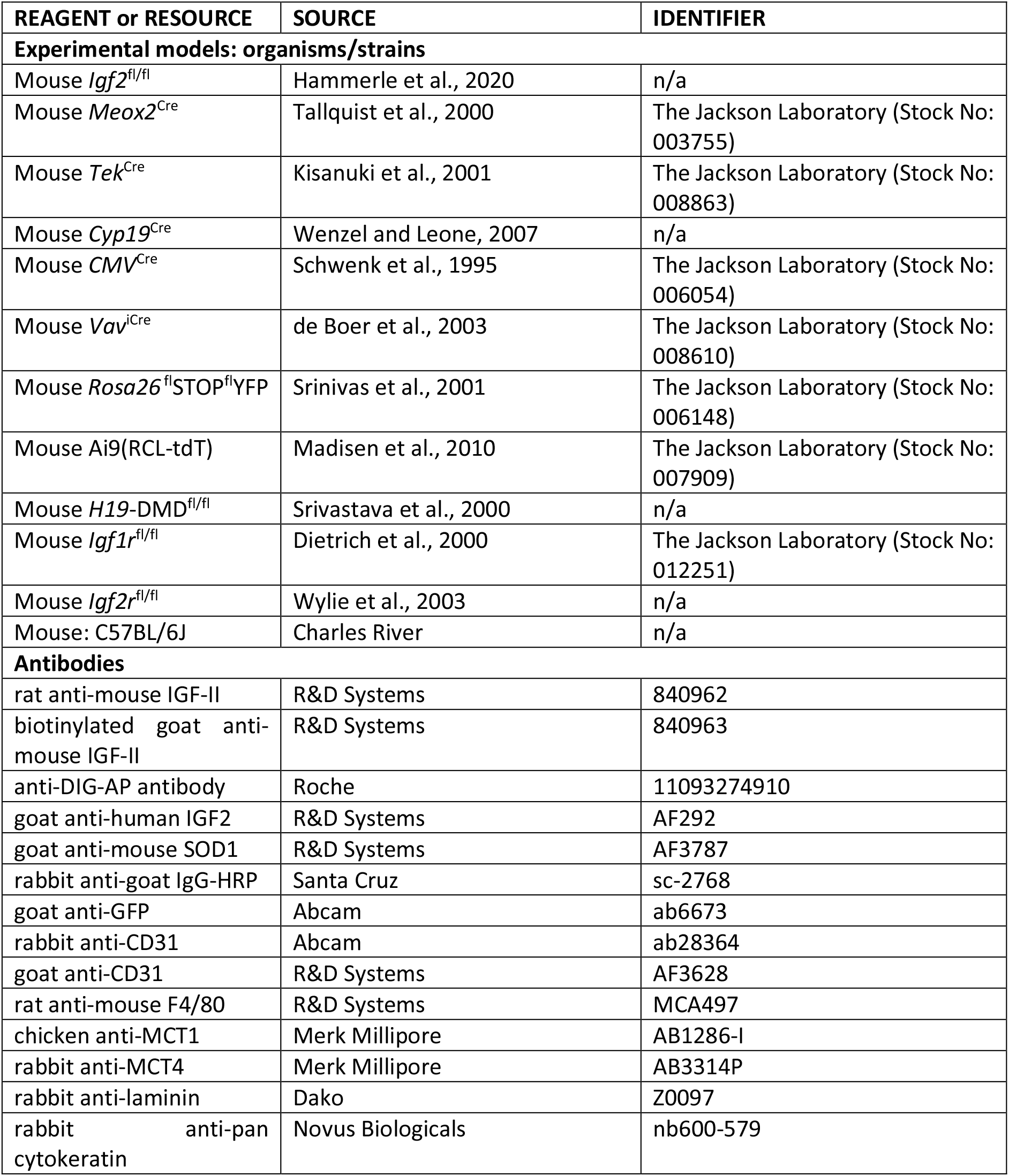
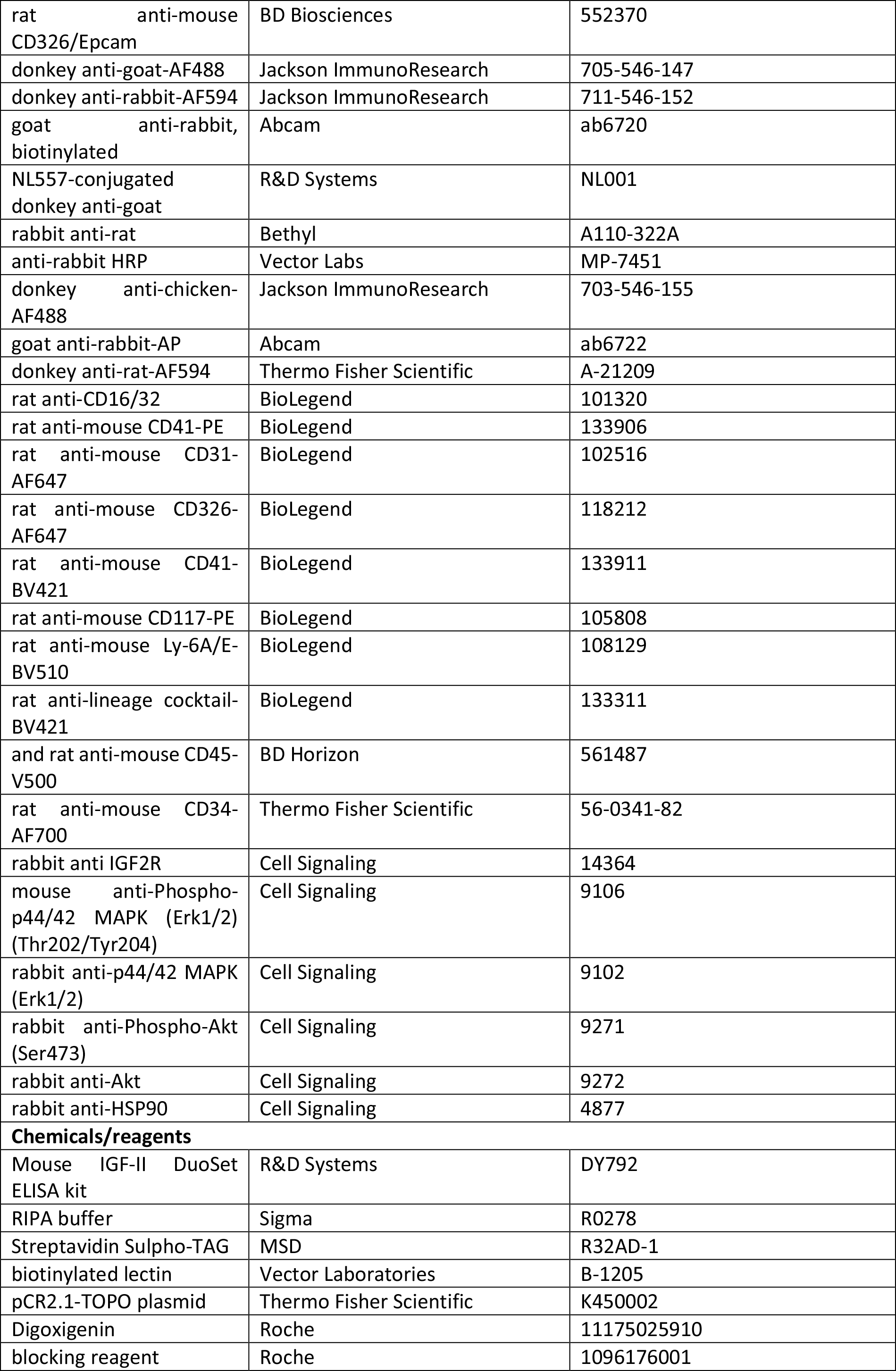
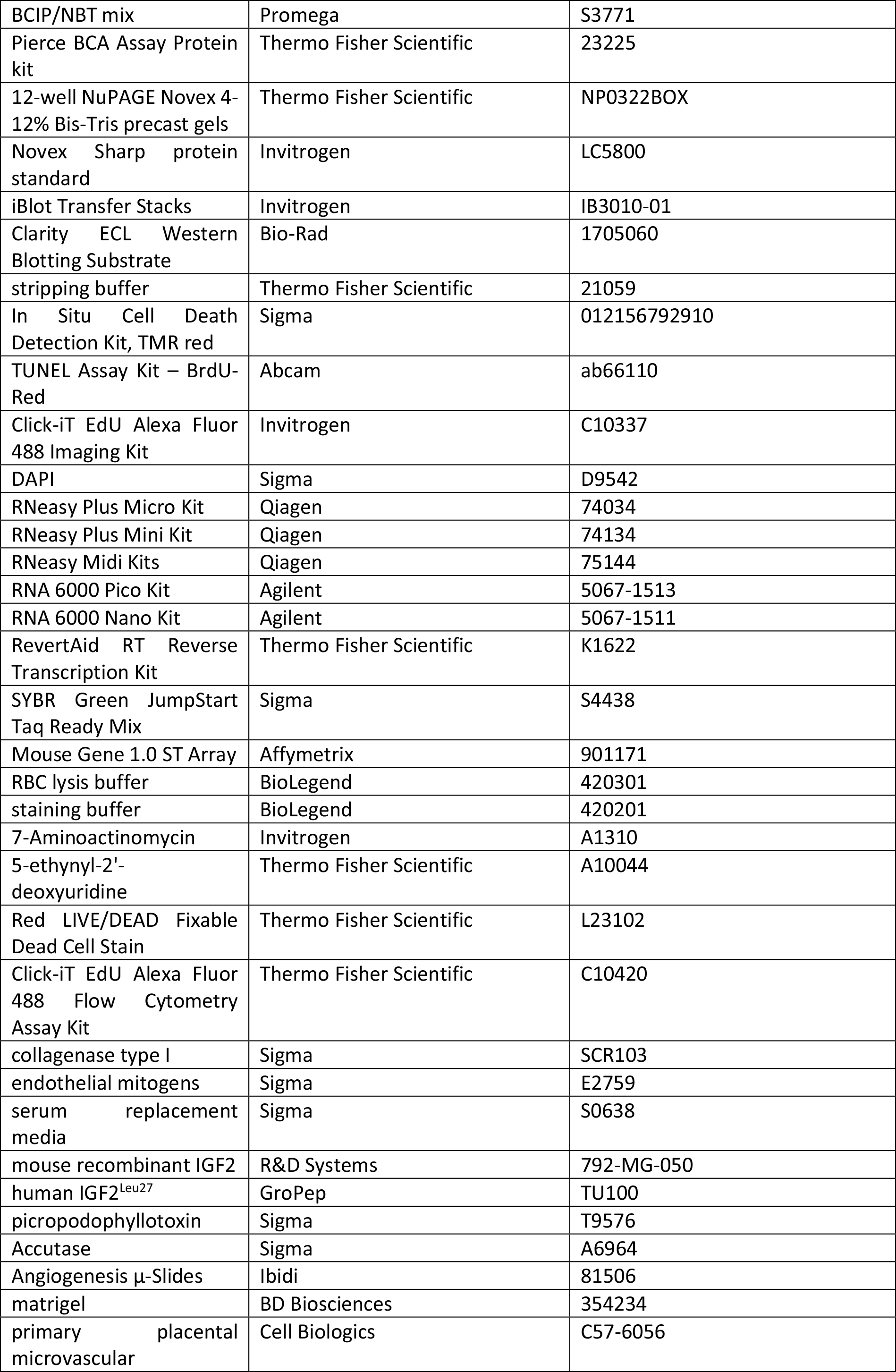
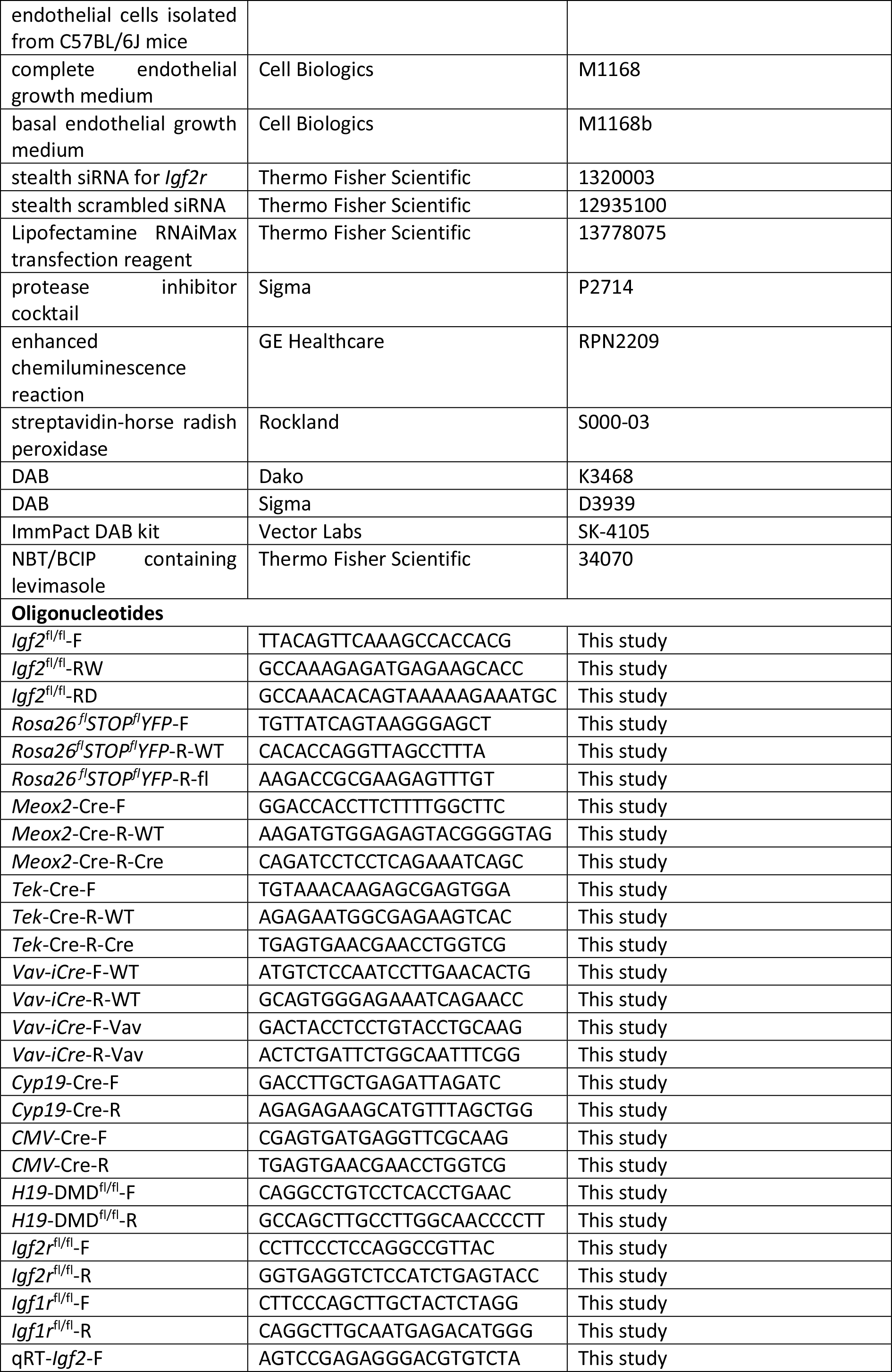
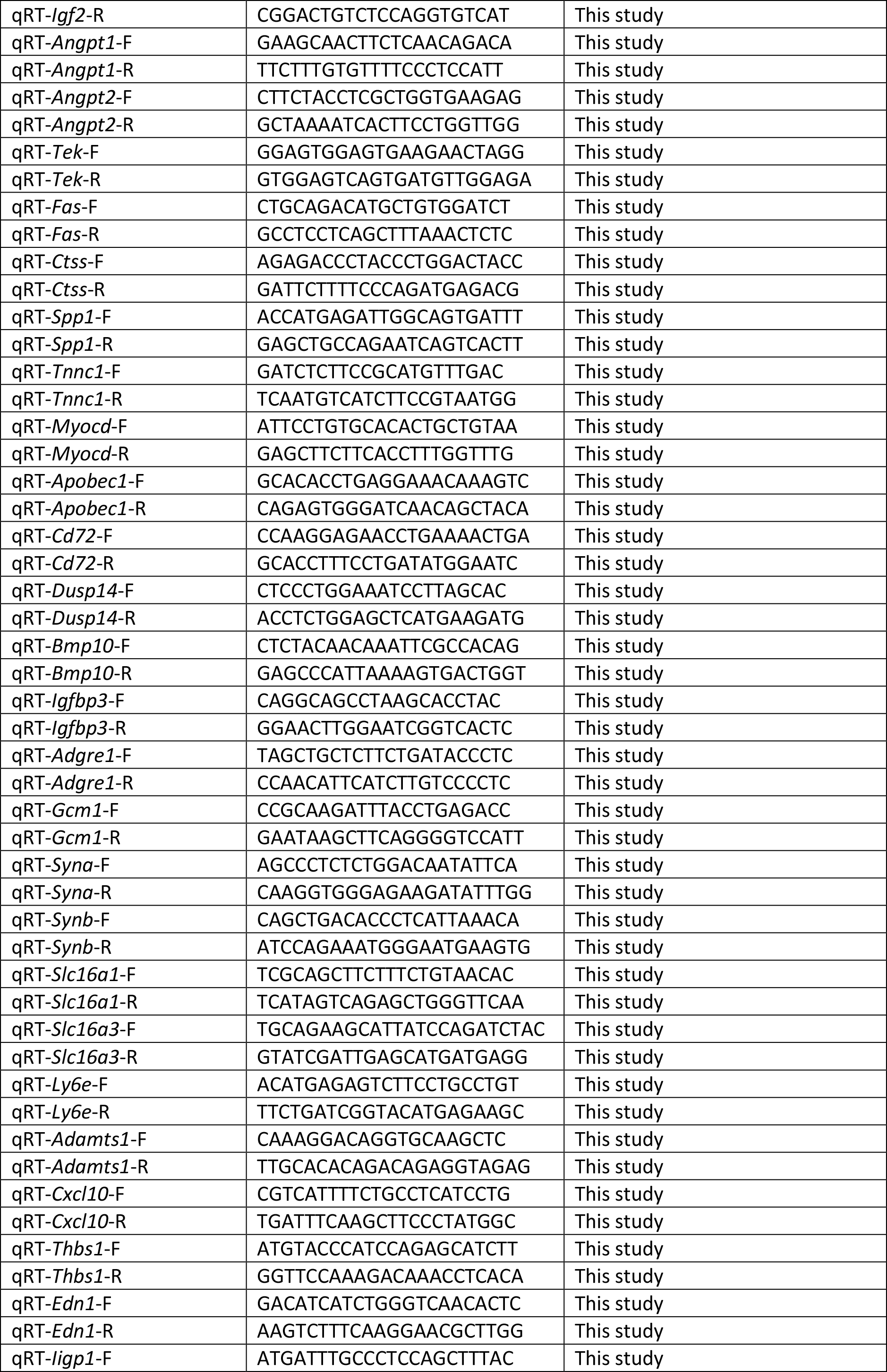
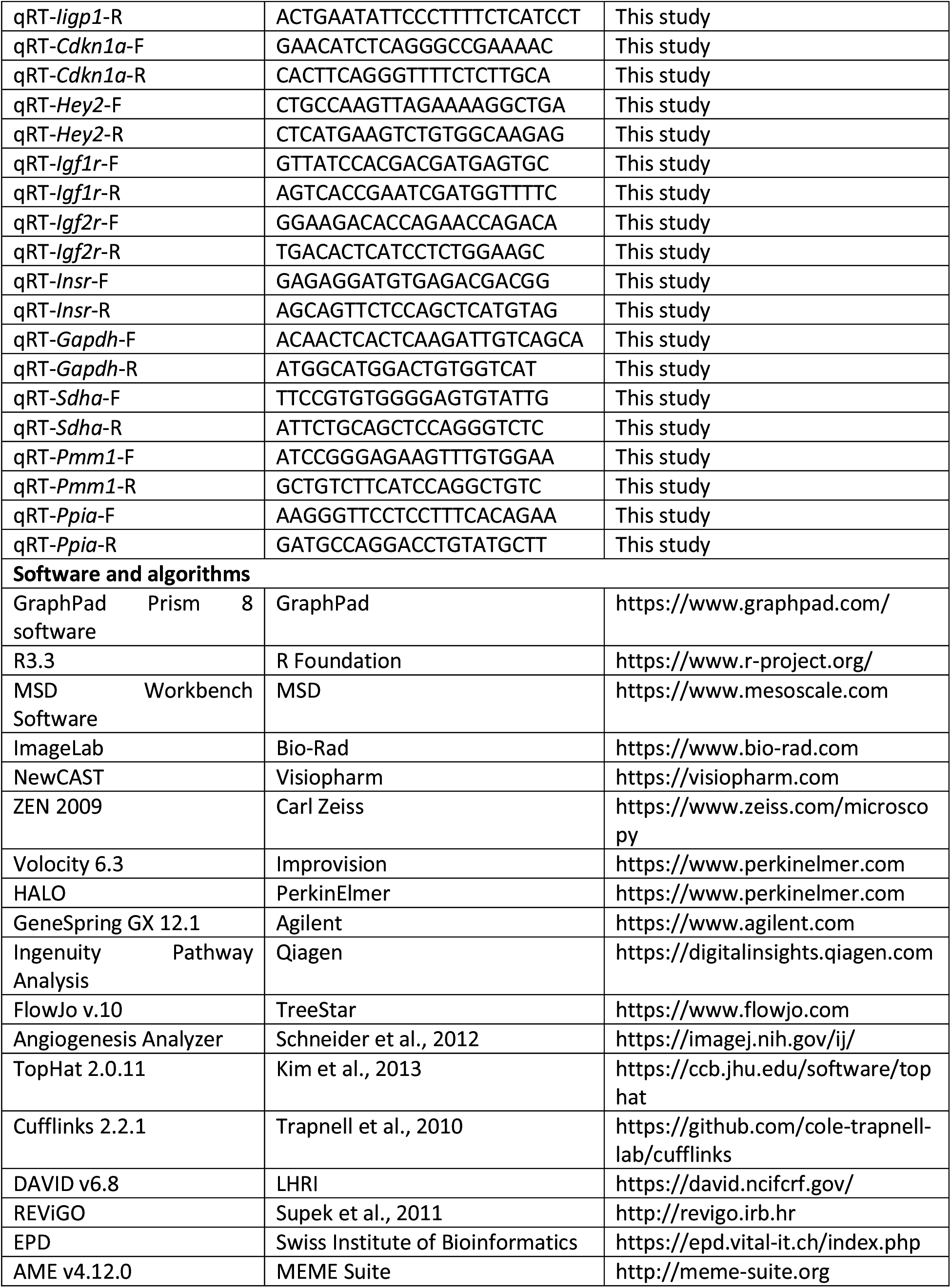

## RESOURCES AVAILABILITY

### Lead contact

Further information and requests for resources and reagents should be directed to and will be fulfilled by the Lead Contact, Miguel Constância.

### Materials availability

This study did not generate new unique reagents.

### Data and code availability

Processed gene expression data from expression microarray and RNA-seq comparisons are available in Tables S1–S3 and the corresponding raw data have been deposited in the Gene Expression Omnibus (GEO) under the accession numbers GSE125434 and GSE179549. Other data and materials are available upon request from the corresponding authors.

## EXPERIMENTAL MODEL AND SUBJECT DETAILS

### Mice

Mice were bred, maintained and mated under pathogen-free conditions at the University of Cambridge Phenomics Unit (West Forvie), in accordance with the University of Cambridge Animal Welfare and Ethical Review Body and the United Kingdom Home Office Regulations. The morning of the copulation plug discovery was counted as embryonic day 1 (E1).

The *Igf2*^fl/fl^ mice were generated in our laboratory (Hammerle et al., 2020). *Meox2*^Cre^ mice (Tallquist et al., 2000), *Tek*^Cre^ mice (Kisanuki et al., 2001), *CMV*^Cre^ mice (Schwenk et al., 1995) and *Igf1r*^fl/fl^ mice (Dietrich et al., 2000) were imported from the Jackson Laboratory (Maine, USA). *Meox2*^Cre^ is active starting at E5 in the epiblast, which gives rise to the entire embryo proper and FPEC (Tallquist et al., 2000). *Tek*^Cre^ (also known as *Tie2*^Cre^) activity starts at E7.5 in the endothelial cell lineage, including FPEC (Kisanuki et al., 2001). *CMV*^Cre^ activity starts soon after fertilization and induces ubiquitous deletion of floxed alleles in all tissues, including the germline (Schwenk et al., 1995). *Cyp19*^Cre^ mice (Wenzel and Leone, 2007) were kindly provided by Prof. Gustavo Leone (Medical University of South Carolina). *Cyp19*^Cre^ is active from E6.5 in the early diploid trophoblast cells that give rise to spongiotrophoblast, giant cells, and labyrinthine trophoblast cells (Wenzel and Leone, 2007). *Vav*^iCre^ mice, in which expression of an optimized variant of Cre is expressed in all hematopoietic cells but not in endothelial cells (de Boer et al., 2003), were kindly provided by Dr. Bidesh Mahata (University of Cambridge). (*Rosa26* ^fl^STOP^fl^YFP mice (Srinivas et al., 2001) were kindly provided by Dr. Martin Turner (The Babraham Institute, Cambridge), Ai9(RCL-tdT) mice (Madisen et al., 2010) by Prof. William Colledge (University of Cambridge), *H19*-DMD^fl/fl^ mice (Srivastava et al., 2000) and *Igf2r*^fl/fl^ mice (Wylie et al., 2003) by Prof. Bass Hassan (University of Oxford). Deletion of *H19*-DMD leads to reactivation of the silent maternal *Igf2* allele, as well as down-regulation of *H19* mRNA levels (Srivastava et al., 2000).

All strains were bred into an inbred C57BL/6J genetic background for >10 generations, with the exception of *Vav*^iCre^ strain that was maintained on an inbred C57BL/6N genetic background. For all crosses (Table S5), the parent transmitting the floxed allele was also homozygous for the *Rosa26* ^fl^STOP^fl^YFP allele. Thus, YFP expression provided an internal control for efficiency of Cre deletion (see Figures S1, S3, S6 and S7). For all crosses fl/+ and +/fl as superscripts mean that the offspring has inherited the floxed allele from the mother and father, respectively; Cre/+ and +/Cre as superscripts mean that the offspring has inherited the Cre recombinase from the mother and father, respectively; combination of fl/+, +/Cre means deletion of maternal floxed allele and combination of +/fl, Cre/+ means deletion of paternal floxed allele (see Table S5). Genotyping was performed by standard PCR using DNA extracted from ear biopsies (adult mice) or tail DNA (fetuses). PCR was performed using the Red Taq Ready PCR system (Sigma) (see list of primers in Table S6), followed by separation of PCR amplicons by agarose gel electrophoresis.

## METHOD DETAILS

### Plasma IGF2 measurements

IGF2 measurements were performed with the Mouse IGF-II DuoSet ELISA kit (R&D Systems – DY792), using an assay adapted for the MesoScale Discovery electrochemiluminescence immunoassay platform (MSD). Briefly, MSD standard-bind microtitre plates were first coated with 30µl capture antibody (Rat Anti-Mouse IGF-II, R&D Systems – 840962) diluted to 7.2 µg/ml in PBS, sealed, and incubated overnight at 4°C. After three washes with MSD wash (0.1% Tween 20 in PBS), the plates were loaded with 20µl ELISA Diluent RD5-38 per well, plus 10µl standard or plasma (diluted 50 fold in RIPA buffer, Sigma – R0278). The plates were then sealed and incubated for two hours at room temperature on a plate shaker. After three washes with MSD wash, the wells were plated with 25µl detection antibody (Biotinylated Goat Anti-Mouse IGF-II, R&D Systems – 840963), diluted to 0.72 µg/ml in PBS, sealed, and incubated for one hour at room temperature on a plate shaker. Following three additional washes with MSD wash, the wells were plated with 25µl MesoScale Discovery Streptavidin Sulpho-TAG (MSD – R32AD-1), diluted 1:1000 in the MSD Diluent 100, sealed and incubated for 30 minutes at room temperature on a plate shaker. After three final washes with MSD wash, the wells were plated with 150µl of MSD Read Buffer T (1x) and the reading was performed on the MSD s600 analyser. Each sample was measured in duplicate and the results were calculated against the standard curve, using the MSD Workbench Software.

### *Igf2* mRNA *in situ* hybridization

*In situ* hybridization was performed as described (Simmons et al., 2008), with minor modifications. Briefly, a region of 415bp spanning *Igf2* coding exons 4-6 was PCR amplified using primers: 5’-CACGCTTCAGTTTGTCTGTTCG-3’ and 5’-GCTGGACATCTCCGAAGAGG-3’ and E14 placental cDNA as template. The PCR amplicon was cloned into a pCR2.1-TOPO plasmid (Thermo Fisher Scientific – K450002). Sense (S) and antisense (AS) RNA probes were generated and labelled with Digoxigenin (DIG) by *in vitro* reverse transcription, according to manufacturer’s instructions (Roche – 11175025910). E14 fetuses and placentae were collected in ice-cold PBS and fixed overnight in 4% paraformaldehyde in 0.1% diethylpyrocarbonate (DEPC)-PBS at 4°C. Tissues were then dehydrated and embedded in paraffin, using RNase-free conditions. Tissue sections (7μm thick) mounted on polysine slides were de-waxed, rehydrated in PBS, post-fixed in 4% paraformaldehyde for 10 minutes, digested with proteinase K (30μg/ml) for 10 min at room temperature, acetylated for 10 minutes (acetic anhydride, 0.25%) and hybridized overnight at 65°C in a humidified chamber with DIG-labeled probes diluted in hybridization buffer. Two 65°C post-hybridization washes (1×SSC, 50% formamide, 0.1% tween-20) followed by two room temperature washes in 1×MABT were followed by 30 minutes RNAse treatment. Sections were blocked for 1 hour in 1×MABT, 2% blocking reagent (Roche – 1096176001), 20% heat-inactivated goat serum and then incubated overnight with anti-DIG-AP antibody (Roche – 11093274910; 1:2,500 dilution) at 4°C. After 4x20 min washes in 1×MABT, slides were rinsed in 1×NTMT and incubated with BCIP/NBT mix in NTMT buffer, according to manufacturer’s instructions (Promega – S3771). Slides were counterstained with nuclear fast red, dehydrated, cleared in xylene and mounted in DPX mounting medium. Pictures were taken with an Olympus DP71 bright-field microscope fitted with a camera.

### Western blot analysis

Tissues were lysed in ∼10μl/mg tissue RIPA buffer (Sigma – R0278), then the lysates were spun at 3,000 RPM and 4°C for 15 minutes. The supernatants were transferred into new tubes and protein concentrations were quantified using the Pierce BCA Assay Protein kit (Thermo Fisher Scientific – 23225). 60μg total protein were mixed with SDS gel loading buffer, then denatured at 70°C for 10 minutes and loaded into 12-well NuPAGE Novex 4-12% Bis-Tris precast gels (Thermo Fisher Scientific – NP0322BOX). The pre-stained Novex Sharp protein standard (Invitrogen – LC5800) was used as protein marker. After electrophoresis for 40 minutes at 200V and 4°C, the proteins were transferred onto nitrocellulose membranes, using the iBlot Transfer Stacks (Invitrogen – IB3010-01) and the iBlot Gel Transfer Device set for 7 minutes at 20V. Blocking was performed for one hour at 4°C in 5% semi-skimmed milk (Marvel) dissolved in TBS-T. The membranes were then incubated overnight at 4°C with the primary antibody dissolved in 0.5% milk in TBS-T (goat anti-human IGF2, 1:1,000, R&D Systems – AF292-NA or goat anti-mouse SOD1, 1:50,000, R&D Systems – AF3787). After 2x10 minutes washes with milliQ water and 2x10 minutes washes with TBS-T, the blots were incubated for one hour at room temperature with the secondary antibody dissolved in TBS-T containing 3% semi-skimmed milk (rabbit anti-goat IgG-HRP, 1:2,500, Santa Cruz sc-2768). The blots were then washed as above, exposed to substrate (Clarity ECL Western Blotting Substrate, Bio-Rad – 1705060) for 5 minutes and imaged with the Bio-Rad GelDoc system. Stripping of antibodies was carried out using a stripping buffer (Thermo Fisher Scientific – 21059) for 15 minutes at room temperature. The band intensities were quantified using the ImageLab software (Bio-Rad) and expressed as IGF2/SOD1 ratios.

### Placenta stereology

Placenta stereology analyses for the *Igf2*^EpiKO^ and *Igf2*^ECKO^ models were performed as described (Coan et al., 2004) in placentae (n=5–7) collected from three litters at each developmental stage. Briefly, the placentae were weighted, then halved and each half placenta weighted again. A half was fixed in 4% paraformaldehyde in PBS at 4°C overnight, then dehydrated and embedded in paraffin wax. The paraffin blocks were exhaustively sectioned using a microtome at 7μm thickness. Placental sections spaced 140 μm apart were hematoxylin-eosin stained and stereological measurements of placental layers were done using the NewCAST system (Visiopharm, Hoersholm, Denmark), using the point counting method (Coan et al., 2004).

The corresponding placental halves were fixed for 6 hours with 4% glutaraldehyde in 0.1 M PIPES buffer, washed with 0.1 M PIPES buffer, and treated with 1% osmium tetroxide. The samples were then resin-embedded and 1μm thick sections, obtained close to the placental midline, were stained with methylene blue. Analysis of Lz components was done using the NewCAST system (Visiopharm) with meander sampling of ∼25% of the Lz area.

Placental stereology for *Igf2*^HCKO^ model was performed as described (De Clercq et al., 2020). Briefly, placental samples were embedded in paraffin as described above, sectioned and then double-labelled for lectin and cytokeratin, which allows the identification of Lz constituents. The proportion of FC, MBS and LT was quantified using the NewCAST system and the point counting method, as described above.

### Transmission electron microscopy

Analysis of E16 *Igf2*^EpiKO^ mutant and control placentae by transmission electron microscopy was performed as previously described (Coan et al., 2005). Briefly, resin-embedded 1 μm thick sections, cut near placental midline and stained with methylene blue as described in the previous section (placenta stereology) were used to identify regions of interest. Thin sections (50 nm) were stained with uranyl acetate and lead citrate, and viewed using a Philips CM100 transmission electron microscope at 80 kV.

### Immunostainings

Immunohistochemistry or immunofluorescence conditions are listed in Table S7. TUNEL staining was performed using the In Situ Cell Death Detection Kit, TMR red (Sigma – 012156792910), or the TUNEL Assay Kit – BrdU-Red (Abcam – ab66110) according to manufacturer’s protocols. EdU staining was done with the Click-iT EdU Alexa Fluor 488 Imaging Kit (Invitrogen – C10337), according to manufacturer’s instructions. For all immunofluorescence stains, DAPI (Sigma – D9542) was used to label the nuclei. For all immunohistochemistry, images were taken with an Olympus DP71 bright-field microscope. Immunofluorescence image acquisition was performed using a LSM510 Meta confocal laser scanning microscope (Carl Zeiss, Jena, Germany) and the ZEN 2009 software or a SP8 laser-scanning confocal microscope (Leica, Mannheim, Germany). Fluorescence semi-quantification analysis was performed using Volocity 6.3 (Improvision). Counting of TUNEL^+^ and F4/80^+^ cells was performed using HALO image analysis software (PerkinElmer).

### qRT-PCR analysis

Total RNA was extracted using RNeasy Plus Kits (Qiagen – 74134 and 74034). RNA concentration was measured by NanoDrop (Thermo Fisher Scientific) and quality was assessed in agarose gels. RNA extracted from FACS isolated cells was quantified and assessed for quality using the RNA 6000 Pico Kit (Agilent – 5067-1513) and an Agilent 2100 Bioanalyzer. Reverse transcription was performed using the RevertAid RT Reverse Transcription Kit (Thermo Fisher Scientific – K1622). qRT-PCR was performed with the SYBR Green JumpStart Taq Ready Mix (Sigma – S4438) and custom-made primers (Table S6) using an ABI Prism 7900 system (Applied Biosystems). For gene expression normalization, we used four housekeeping genes (*Gapdh*, *Sdha*, *Pmm1*, *Ppia*). Levels of expression were calculated using the 2^-ΔΔCt^ method (Livak et al., 2001).

### Expression microarray analysis

Total RNA was extracted from E19 male placental Lz using RNeasy Midi Kits (Qiagen – 75144) and quantity and quality were verified using RNA 6000 Nano Kit (Agilent – 5067-1511) and an Agilent 2100 Bioanalyzer. Only RNA samples with RNA integrity numbers (RIN) >9.0 were used. Array profiling was performed using the Mouse Gene 1.0 ST Array (Affymetrix – 901171) and the analysis of the data was performed using GeneSpring GX 12.1 (Agilent, Santa Clara, CA, USA), with two algorithms: RMA (Robust Multiarray Average) and PLIER (Probe Logarithmic Intensity Error). Only genes with log_2_ fold change > 0.3 predicted by both algorithms were listed as DEGs. Pathway analysis was performed using Ingenuity Pathway Analysis (version 2012).

### Flow cytometry analyses

For flow cytometry analyses of FPEC, placental Lz samples were micro-dissected in ice-cold PBS. Tissue dissociation into single cells was achieved by digestion at 37°C for 45 minutes with a 0.1% collagenase P solution, aided by mechanical dissociation with needles of decreasing diameter. The cells were then passed through 70-µm cell strainers and washed once in ice-cold PBS + 0.1% BSA. Erythrocytes were lysed using the RBC lysis buffer (BioLegend – 420301). Pelleted cells were then re-suspended in 100µl staining buffer (BioLegend – 420201), counted using the Cedex XS Analyser (Roche) and diluted at 1,000 cells/µl. Blocking of Fc receptors was performed by incubation at 4°C for 20 minutes with an unlabelled anti-CD16/32 (1 μg/million cells; BioLegend – 101320). The cells were then incubated for one hour at 4°C in the dark with a mix of rat anti-mouse CD41 (labelled with Phycoerythrin, PE) (BioLegend – 133906; 0.25 µg per million cells), rat anti-mouse CD31 (labelled with AF647) (BioLegend – 102516; 0.25 µg per million cells) and rat anti-mouse CD45 (labelled with V500) (BD Horizon – 561487; 0.4 µg per million cells) in 200µl staining buffer. Stained cells were washed twice in 1ml staining buffer, re-suspended in PBS containing a viability marker (7-AAD – 7-Aminoactinomycin, Invitrogen – A1310), filtered again through 70-µm cell strainers and incubated on ice for 5 minutes. Flow cytometry analysis was performed with a BD FACSCantoII machine (BD Biosciences) and 100,000 events were recorded for each sample. FSC files were analysed with the FlowJo_V10 software, using single-cell discrimination and gating based on single-stained controls. FPEC were identified as 7AAD^-^ /CD31^+^/CD41^-^ cells.

For flow cytometry analyses of EPCAM^high^ positive cells, whole E12 placentae were dissociated into single cells as described above. After erythrocyte lysis, cell counting and blocking of Fc receptors using an unlabelled anti-CD16/32 antibody, cells were incubated for one hour at 4°C in the dark with AF647 rat anti-mouse CD326 (Epcam) antibody (BioLegend – 118212; 0.25 µg per million cells) in 200µl staining buffer. Stained cells were washed as above, incubated with the viability marker 7-AAD and filtered through 70-µm cell strainers. Flow cytometry analysis was performed with a BD FACSCantoII machine (BD Biosciences) and FSC files were analysed with the FlowJo_V10 software, using single-cell discrimination and gating based on single-stained controls.

### Flow cytometry analysis of FPEC proliferation

Pregnant female mice received intraperitoneal (i.p.) injections with 50µg of 5-ethynyl-2’-deoxyuridine (EdU)/g body weight (Thermo Fisher Scientific – A10044), 16 hours prior to tissue collection. Placental Lz dissociation into single cells was performed as above. Cells re-suspended at a concentration of 1000 cells/µl were incubated for 30 minutes at 4°C with 1 µl Red LIVE/DEAD Fixable Dead Cell Stain (Thermo Fisher Scientific – L23102). After one wash in PBS, the cells were pre-incubated for 20 minutes at 4°C in the dark with unlabelled rat anti-mouse CD16/32 (BioLegend – 101320, 1 μg/million cells), then for 1 hour at 4°C in the dark with a 1:1 mix of rat anti-mouse CD41 (labelled with BV421) (BioLegend – 133911; 0.25 µg per million cells) and rat anti-mouse CD31 (labelled with AF647) (BioLegend – 102516; 0.25 µg per million cells) in staining buffer. After two washes with staining buffer, the cells were stained using the Click-iT EdU Alexa Fluor 488 Flow Cytometry Assay Kit (Thermo Fisher Scientific – C10420), according to manufacturer’s instructions. Flow cytometry analysis was performed using a BD LSRFortessa cell analyser (BD Biosciences). FSC files were analysed with the FlowJo_V10 software, using single-cell discrimination and gating based on single-stained controls. Proliferating FPEC were identified as viable EdU^+^/CD31^+^/CD41^-^ cells.

### FPEC and HC isolation by FACS

For sorting FPEC, single cell preparation and staining was performed as above. For sorting HC, entire placentae collected at E13 were used for single cell preparation as described above. Single cell suspensions were pre-incubated for 20 minutes at 4°C in the dark with unlabelled rat anti-mouse CD16/32 (BioLegend – 101320, 1 μg/million cells), then stained for one hour at 4°C in the dark with a mix of rat anti-mouse CD117/c-kit (labelled with PE) (BioLegend – 105808; 0.4 µg per million cells), rat anti-mouse CD34 (labelled with AF700) (ThermoFisher Scientific – 56-0341-82; 1 µg per million cells), rat anti-mouse Ly-6A/E (Sca1) (labelled with BV510) (BioLegend – 108129; 5 µl per million cells) and rat anti-lineage cocktail (labelled with BV421) (BioLegend – 133311; 5 µl per million cells) in staining buffer and washed twice. FACS was done using an Aria-Fusion cell sorter (BD Bioscience), with exclusion of cell duplets and dying cells (7AAD^+^). Cell fractions (endothelial, non-endothelial and hematopoietic cells) were then spun at 3,000 RPM and 4°C for 3 min, the excess of sorting liquid was removed and cell pellets were flash frozen in liquid nitrogen and stored at -80°C until used for RNA extraction.

### Primary FPEC isolation, culture and tube formation assay

Primary FPEC were isolated as previously described (Branco-Price et al., 2012) and adapted here to placental Lz (E16). Briefly, placental Lz were micro-dissected on ice in RPMI containing 1% penicillin/streptomycin. All samples from one litter were pooled, minced and digested for 90 minutes at 37°C in 2 mg/ml collagenase type I (Sigma – SCR103) in HBSS containing 2mM CaCl2, 2mM MgSO4, and 20mM HEPES. The digests were filtered through 70μm nylon cell strainers and washed in HBSS. The cell pellets were then resuspended in PBS containing 0.1% BSA and incubated with anti-CD31-coated magnetic beads for one hour at 4°C. Cells coated with beads were cultured in endothelial cell growth medium consisting of low glucose DMEM:F12 with 1% nonessential amino acids, 2mM sodium pyruvate, buffered with 20mM HEPES and supplemented with 20% FBS and 75μg/ml endothelial mitogens (Sigma – E2759). The cells were incubated at 37°C in 5% O2 and 5% CO2. After four days, the dead cells were washed and new media was added, additionally supplemented with 20μg/ml Heparin. Sub-confluent cells (∼80%) at passage one (around 10 days in culture) were washed and then cultured in 5% serum replacement media (Sigma – S0638) for ∼40 hours. From each litter we used cells at passage one for treatment with 50 ng/ml mouse recombinant IGF2 (R&D Systems, 792-MG-050; dissolved in PBS), 1000 ng/ml human IGF2^Leu27^ (GroPep – TU100; dissolved in 10mM HCl), 500nM picropodophyllotoxin (PPP, Sigma – T9576; dissolved in DMSO) or 500nM PPP + 50 ng/ml IGF2, or appropriate vehicle control. The cells were harvested with Accutase (Sigma – A6964) and counted using the ADAM^TM^ Automated cell counter (NanoEnTek Inc) and 3,000 cells were seeded into 15-well Angiogenesis µ-Slides (Ibidi – 81506) preloaded with 10µl matrigel/well (BD Biosciences – 354234). Photographs were taken at 30 min, 4, 6 and 8 hours using an EVOS FL Cell Imaging system (Thermo Fisher Scientific). Each experiment was performed on 5-6 litters for every treatment. For each tube formation assay, we used five wells seeded with primary FPEC exposed to the treatment agent with equivalent numbers of the corresponding vehicle. Quantification of tubular network structures was performed using the Angiogenesis Analyzer software in ImageJ (Schneider et al., 2012).

### siRNA knockdown of *Igf2r*

Small interfering RNA (siRNA) knockdown of *Igf2r* was performed on primary placental microvascular endothelial cells isolated from C57BL/6J mice (Cell Biologics, C57-6056) and grown in Cell Biologics’ complete growth medium (M1168) under standard culture conditions (37°C in 21% O2 and 5% CO2). Endothelial cells were transfected with stealth siRNA for *Igf2r* or scrambled siRNA (Thermo Fisher Scientific 1320003 and 12935100, respectively) using Lipofectamine RNAiMax transfection reagent (Thermo Fisher Scientific 13778075).

The impact of *Igf2r* knockdown on endothelial cell proliferation rates, with or without exogenous IGF2 stimulation, was analysed using a previously described protocol (Woods et al., 2017). In brief, 10,000 endothelial cells transfected with either scrambled or *Igf2r* siRNA were plated in basal medium (M1168b, which does not contain VEGF, ECGS, EGF and FBS) supplemented with hydrocortisone, heparin and serum replacement (Sigma, S0638) in the presence or absence of 50 ng/ml recombinant mouse IGF2 (R&D Systems, 792-MG-050), and collected every 24 hours over a period of four days. The number of viable cells was counted using the Countess 3 Automated Cell Counter (Thermo Fisher Scientific), according to manufacturer’s instructions.

To study the impact of *Igf2r* knockdown on intracellular signalling pathways, following 48h of transfection, cells were starved in the basal medium (M1168b) for 20h and then stimulated with recombinant mouse IGF2 (50 ng/ml) and collected at the specific times (1, 5 and 10 min). Total cell extracts were prepared in radioimmunoprecipitation assay buffer (20 mM Tris-HCl, pH 8.0, 137 mM NaCl, 1 mM MgCl2, 1 mM CaCl2, 10% glycerol, 1% NP-40, 0.5% sodium deoxycholate, 0.1% sodium dodecyl sulphate), containing a protease inhibitor cocktail (Sigma, P2714), and incubated at 4°C for 1 h. Western blotting was performed as previously described (Pérez-García et al., 2014). Blots were probed with the following antibodies: rabbit anti IGF2R (Cell Signaling, 14364), mouse anti-Phospho-p44/42 MAPK (Erk1/2) (Thr202/Tyr204) (Cell Signaling, 9106), rabbit anti-p44/42 MAPK (Erk1/2) (Cell Signaling, 9102), rabbit anti-Phospho-Akt (Ser473) (Cell Signaling, 9271), rabbit anti-Akt (Cell Signaling, 9272), rabbit anti-HSP90 (Cell Signaling, 4877). Horseradish peroxidase-conjugated secondary antibodies were from Bio-Rad. Detection was carried out with enhanced chemiluminescence reaction (GE Healthcare, RPN2209) using standard X-ray films. Band intensities were quantified using ImageJ software.

### RNA-sequencing and data analysis

Total RNA was extracted from sorted FPEC by FACS from E16 male placentae using RNeasy Plus Micro Kits (Qiagen – 74034). Quantity and quality were verified using the RNA 6000 Pico Kit (Agilent – 5067-1513) and an Agilent 2100 Bioanalyzer. Only RNA samples with RNA integrity numbers (RIN) >9.0 were used. Total RNA (2 ng) was whole-transcriptome amplified using the Ovation RNA–Seq System V2 (NuGEN). To prepare the RNA–seq libraries the amplified cDNA (2μg per sample) was fragmented to 200bp using a Bioruptor Sonicator (Diagenode), end repaired and barcoded using the Ovation Rapid DR Library System (NuGEN). The libraries were combined and loaded onto an Illumina HiSeq 2500 system for single-end 50bp sequencing at the Genomics Core Facility, Cambridge Institute, CRUK. The reads were aligned onto the mouse GRCm38 genome using TopHat 2.0.11 (Kim et al., 2013). Gene abundance and differential expression were determined with Cufflinks 2.2.1 (Trapnell et al., 2010) and expressed in fragments per kilobase per million mapped reads (FPKM). The cut off for expression was set at ≥1 FPKM. Genes with a linear fold expression change greater than 1.5 and a Benjamini–Hochberg false discovery rate (FDR) <5% were considered differentially expressed.

Functional analysis was performed using DAVID (Database for Annotation, Visualization and Integrated Discovery; v6.8 http://david.abcc.ncifcrf.gov/). Enriched gene ontology (GO) terms with FDR < 5% were considered significant. These terms were then clustered semantically using REViGO (Reduce and Visualize GO) (Supek et al., 2011), which removes redundancy, and ordered according to the log_10_ *P* values.

To search for enrichment of TF binding sites at the promoters of DEG, we used EPD (Eukaryotic Promoter Database – https://epd.vital-it.ch/index.php) to retrieve the DNA sequences from 1,000bp upstream to 100bp downstream of the transcriptional start site (TSS). These sequences were then analysed using AME (Analysis of Motif Enrichment v4.12.0 – http://meme-suite.org/tools/ame) by selecting *Mus musculus* and HOCOMOCO Mouse (v11 FULL) as motif database. Transcriptional network visualization was performed using the Ingenuity Pathway Analysis tool.

## QUANTIFICATION AND STATISTICAL ANALYSIS

No statistical analysis was used to predetermine sample size. Randomization was not used in our animal studies. Placental stereology and histological EdU analyses were performed blinded to genotype. Statistical analyses for fetus, placenta and Lz growth kinetics were performed in R, using a mixed effects model, with litter as a random effect and genotype, developmental stage and the interaction between genotype and developmental stage as fixed effects. Prior to these analyses, fetal, placental and Lz weights were log transformed. All other statistical analyses were performed using GraphPad Prism 8. Statistical significance between two groups was determined by Mann-Whitney tests or two-tailed unpaired t-tests and statistical significance between multiple groups was performed using one-way ANOVA plus Tukey’s multiple comparisons tests or two-way ANOVA plus Sidak’s multiple comparisons tests, as appropriate. The numbers of samples or litters used for each experiment are indicated in figure legends.

