## Supplementary Data for "The Imprinted *Igf2*-*Igf2r* Axis is Critical for Matching Placental Microvasculature Expansion to Fetal Growth"

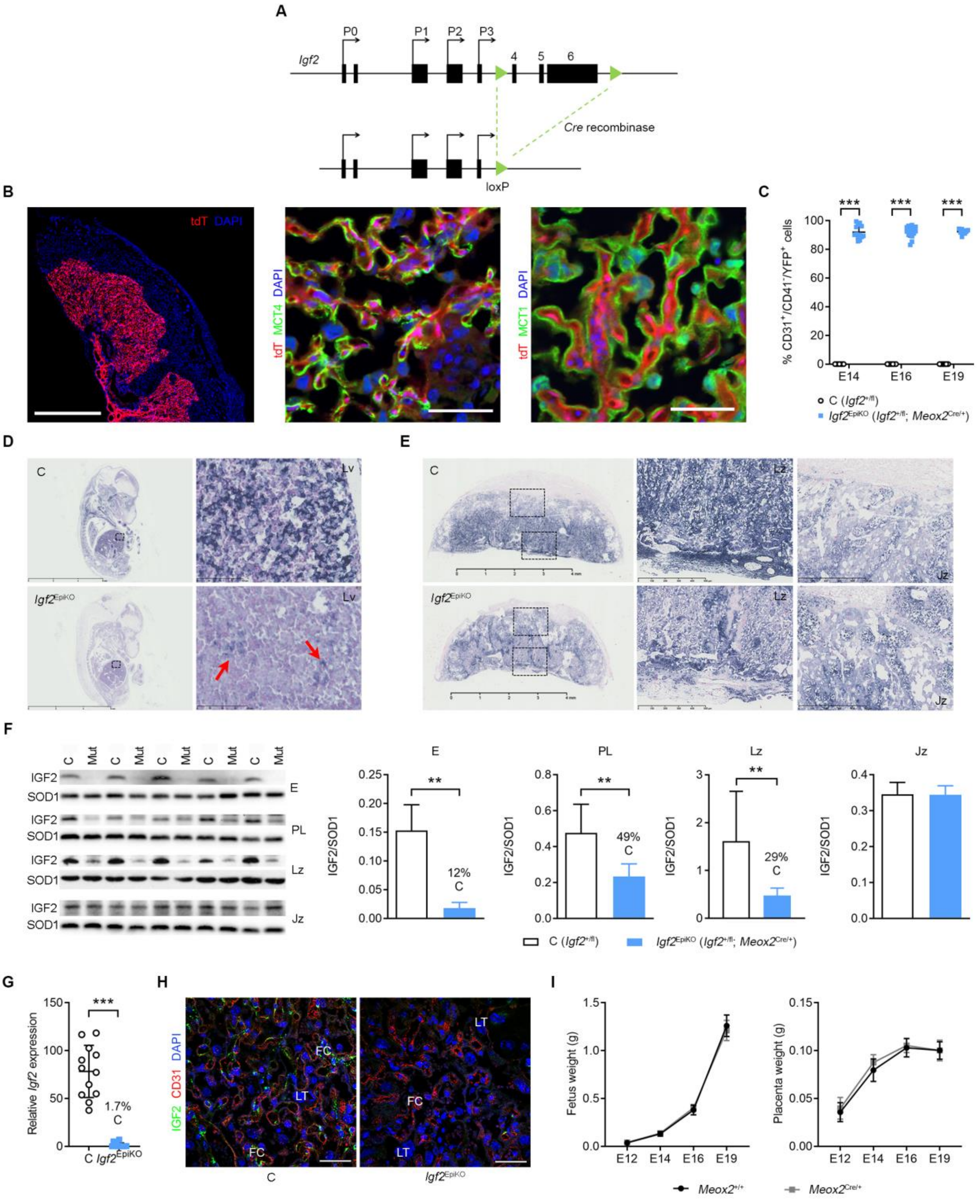


**Figure S1. Specificity and efficiency of *Igf2* deletion in fetal tissues and feto-placental endothelial cells by *Meox2*$^{Cre}$.** (**A**) Schematic representation of the floxed *Igf2* allele. P0-P3 are alternative promoters. Protein-coding exons (4–5), flanked by loxP sites (green triangles), are excised upon Cre-loxP mediated recombination. (**B**) Representative confocal microscopy of a placental frozen section at E16 of gestation, double transgenic for *Meox2*$^{Cre}$ and Ai9(RCL-tdT) reporter. The *Meox2*$^{Cre}$ is not expressed in the syncytiotrophoblast layers, as demonstrated by the lack of immunostaining overlap between the tomato protein (red) and MCT4 (a marker of the syncytiotrophoblast layer II, facing the feto-placental capillaries) or MCT1 (marker of the syncytiotrophoblast layer I, facing the maternal blood spaces). Scale bars are 1 mm (left panel) and 50 µm (middle and right panels). (**C**) Flow cytometry analysis shows that the majority (>80%) of *Igf2*$^{EpiKO}$ mutant FPEC (CD31$^{+}$/CD41$^{-}$ cells) express YFP (activated by *Meox2*$^{Cre}$ mediated deletion of the *Rosa26* $^{fl}$STOP$^{fl}$YFP STOP cassette), thus demonstrating good efficiency of *Meox2*-Cre in these cells (n=9–18 per genotype). (**D**) *Igf2* mRNA in situ hybridization (ISH) in E14 control and mutant fetuses. Dark blue indicates *Igf2* mRNA, with nuclei marked in red. Insets illustrate efficient *Igf2* deletion in the liver (Lv); arrows – small pockets of cells with incomplete *Igf2* deletion (mosaic activity of *Meox2*$^{Cre}$). Scale bars are 6 mm (left) and 100 µm (right). (**E**) *Igf2* mRNA ISH in E14 control and mutant placentae. Insets show reduced *Igf2* mRNA signal in the placental labyrinthine zone (Lz) of mutants, due to its deletion from FPEC, while *Igf2* expression is unchanged in the junctional zone (Jz). Scale bars are 4 mm (left) and 500 µm (right). (**F**) Western blot analysis of pro-IGF2 (18kDa) in cell lysates from whole fetuses (F), whole placenta (PL) micro-dissected placental labyrinthine (Lz) and junctional zones (Jz) at E14, and corresponding data quantification shown as graphs (n=5 per genotype). SOD1 (19 kDa) was used as loading control. (**G**) Efficiency of *Igf2* deletion evaluated by qRT-PCR in fluorescence-activated sorted FPEC (n=12 per genotype). (**H**) Representative immunofluorescence confocal microscopy at E16 showing near complete absence of IGF2 protein within CD31+ feto-placental endothelial cells in *Igf2*$^{EpiKO}$ mutants compared to littermate controls. FC – fetal capillaries, LT – labyrinthine trophoblast. Scale bars are 50µm. (**I**) Fetal and placental growth kinetics are not altered in *Meox2*$^{Cre/+}$ carriers (maternal inheritance) (n=8–30 conceptuses per genotype at each developmental stage). For all graphs, data is shown as individual values or averages ± SD; ** $P < 0.01$, *** $P < 0.001$ calculated by two-way ANOVA plus Sidak's multiple comparisons tests in (C) or Mann Whitney tests in (F) and (G).

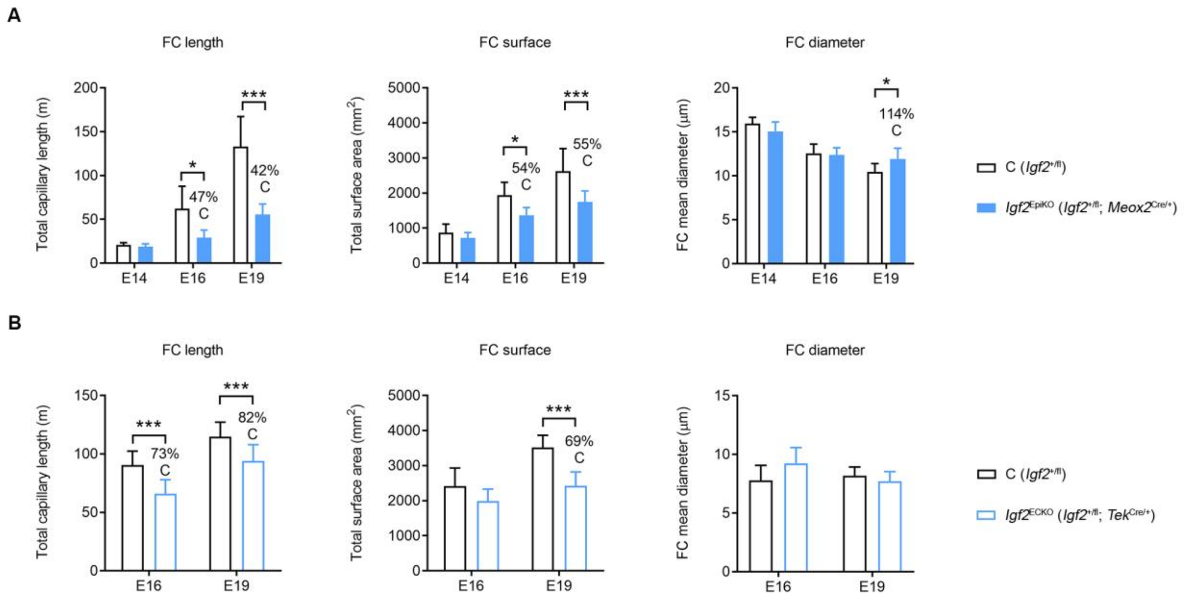


**Figure S2. Impact of *Igf2*$^{\mathbf{EpiKO}}$ and *Igf2*$^{\mathbf{ECKO}}$ deletions on feto-placental capillary expansion during gestation.** (**A**) Parameters of feto-placental capillaries (FC) measured by stereology in *Igf2*$^{\mathrm{EpiKO}}$ mutant (*Igf2*$^{+/\mathrm{fl}}$; *Meox2*$^{\mathrm{Cre}/+}$) versus control (C – *Igf2* $^{+/\mathrm{fl}}$) placentae (n=6 per genotype at each developmental stage). (**B**) Parameters of feto-placental capillaries (FC) measured by stereology in *Igf2*$^{\mathrm{ECKO}}$ mutant (*Igf2*$^{+/\mathrm{fl}}$; *Tek*$^{\mathrm{Cre}/+}$) versus control (C – *Igf2* $^{+/\mathrm{fl}}$) placentae (n=5–7 per genotype at each developmental stage). For all graphs, data is shown as averages ± SD; * $P < 0.05$, *** $P < 0.001$ calculated by two-way ANOVA plus Sidak's multiple comparisons tests.

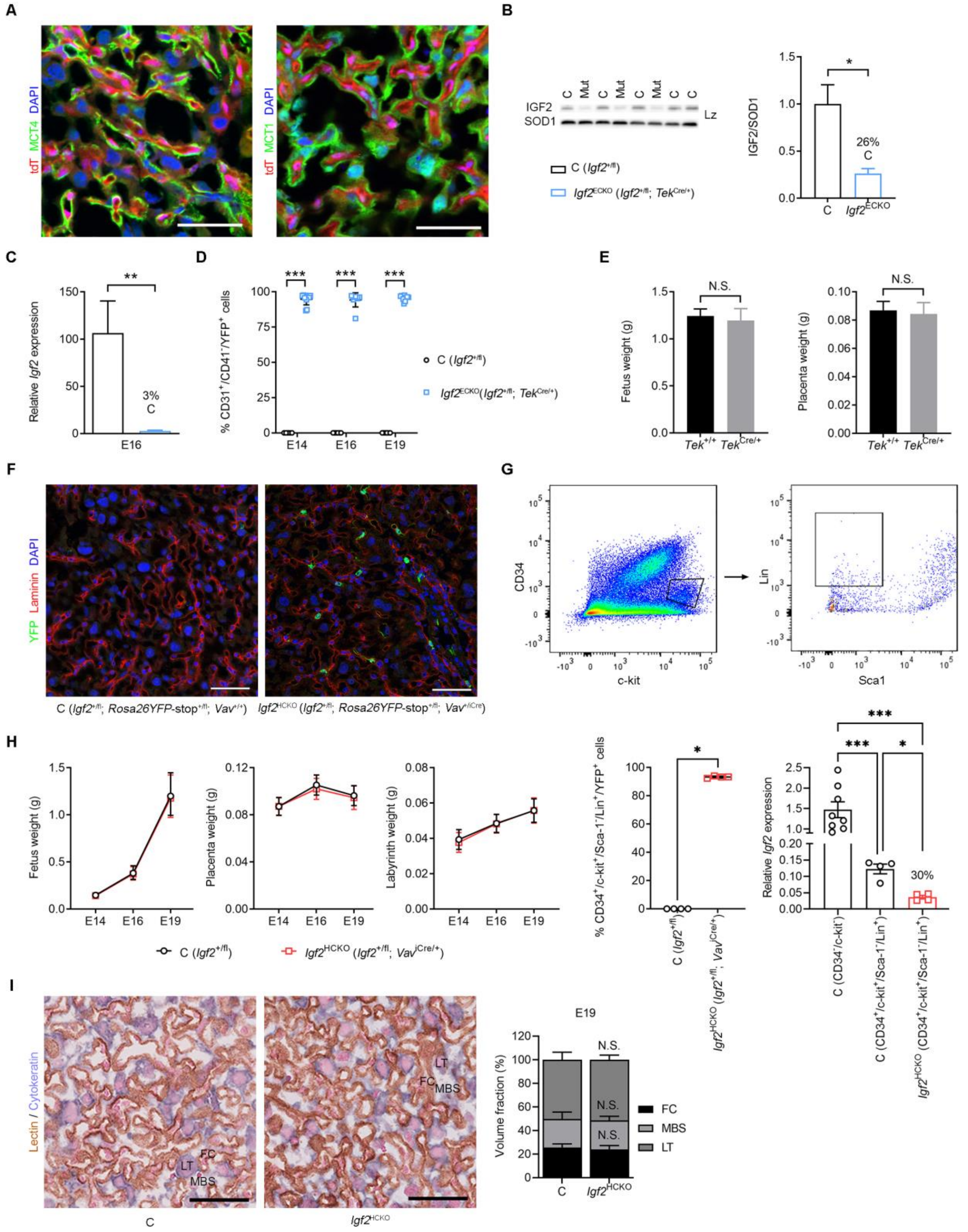


**Figure S3. Specificity and efficiency of *Igf2* deletion in the endothelium by *Tek*$^{Cre}$ and in the hematopoietic lineage by *Vav*$^{iCre}$.** (**A**) Representative confocal microscopy of frozen placental sections from a double transgenic for *TeK*$^{Cre}$ and Ai9(RCL-tdT) reporter at E16 of gestation. The *TeK*$^{Cre}$ is not expressed in the syncytiotrophoblast layers, as demonstrated by the lack of immunostaining overlap between the tomato protein (red) and MCT4 (Syn-TII layer) or MCT1 (Syn-TI layer). Scale bars are 50 µm. (**B**) Western blot analysis of pro-IGF2 (18 kDa) in cell lysates from placental Lz micro-dissected at E16 and corresponding data quantification (n=3 per genotype). SOD1 (19 kDa) was used as internal control for loading. (**C**) Efficiency of *Igf2* deletion evaluated by qRT-PCR in fluorescence-activated sorted FPEC (n=5–7 per genotype). (**D**) Flow cytometry analysis shows that the majority (>80%) of *Igf2*$^{ECKO}$ mutant FPEC express YFP, thus demonstrating good efficiency of *TeK*$^{Cre}$ in these cells (n=5–11 per genotype). (**E**) Fetal and placental growth kinetics are not altered in *TeK*$^{Cre/+}$ carriers (maternal inheritance) at E19 (n=13–15 conceptuses per genotype from 4 independent litters). (**F**) Representative double immunostainings for laminin (marking feto-placental capillaries) and YFP in E19 *Igf2*$^{HCKO}$ mutant and littermate control (C) placentae. YFP expression is activated in cells of hematopoietic lineage by deletion of the floxed STOP cassette within the *Rosa26*$^{fl}$STOP$^{fl}$YFP reporter construct. Scale bars are 50µm. (**G**) Gating strategy used to isolate cells of hematopoietic lineage from E13 placentae by FACS (top) and efficiency of *Igf2* deletion evaluated by qRT-PCR (bottom). The CD34$^{+}$/c-kit$^{+}$ population gated in the top left panel was subsequently used to isolate Lin$^{+}$/Sca1$^{-}$ hematopoietic cells (top right panel). Bottom left: more than 90% of CD34$^{+}$/c-kit$^{+}$/Lin$^{+}$/Sca1$^{-}$ cells are positive for YFP, indicating efficient activity of *Vav*$^{iCre}$ in the hematopoietic cells. Bottom right: measurement of mRNA levels at E13 by qRT-PCR indicates overall low levels of *Igf2* expression in the hematopoietic lineage (approximately one order of magnitude lower than in CD34$^{-}$/c-kit$^{-}$ cells), as well as efficient deletion of *Igf2* in hematopoietic cells of *Igf2*$^{HCKO}$ mutants compared to littermate controls (n=4–8/group). (**H**) Fetal, placental, and placental Lz growth kinetics are not altered in *Igf2*$^{HCKO}$ mutants compared to controls (n=11–25 conceptuses from n=5–6 litters for each developmental stage). (**I**) Left: representative double immunostaining for lectin (brown, marking feto-placental capillaries) and cytokeratin (blue, marking the labyrinthine trophoblast) in E19 placental sections of *Igf2*$^{HCKO}$ mutants and littermate controls. Right: the relative volume fractions occupied by the three main labyrinthine constituents (FC – fetal capillaries, MBS – maternal blood spaces and LT – labyrinthine trophoblast) are not altered in *Igf2*$^{HCKO}$ mutants compared to controls at E19 (n=4–7 samples/group). For all graphs, data is shown as individual values or averages ± SD in (B), (D), (E), (G) – bottom left and (I), SEM in (C) and (G) – bottom right, or 95% confidence intervals (95%CI) in (H); N.S. – statistically non-significant; * $P < 0.05$, ** $P < 0.01$ and *** $P < 0.001$ calculated by Mann Whitney tests in (B), (C), (E) and (G) – bottom left, two-way ANOVA plus Sidak's multiple comparisons tests in (D) and (I), Brown-Forsythe and Welch ANOVA tests in (G) – bottom right and mixed effects model in (H).

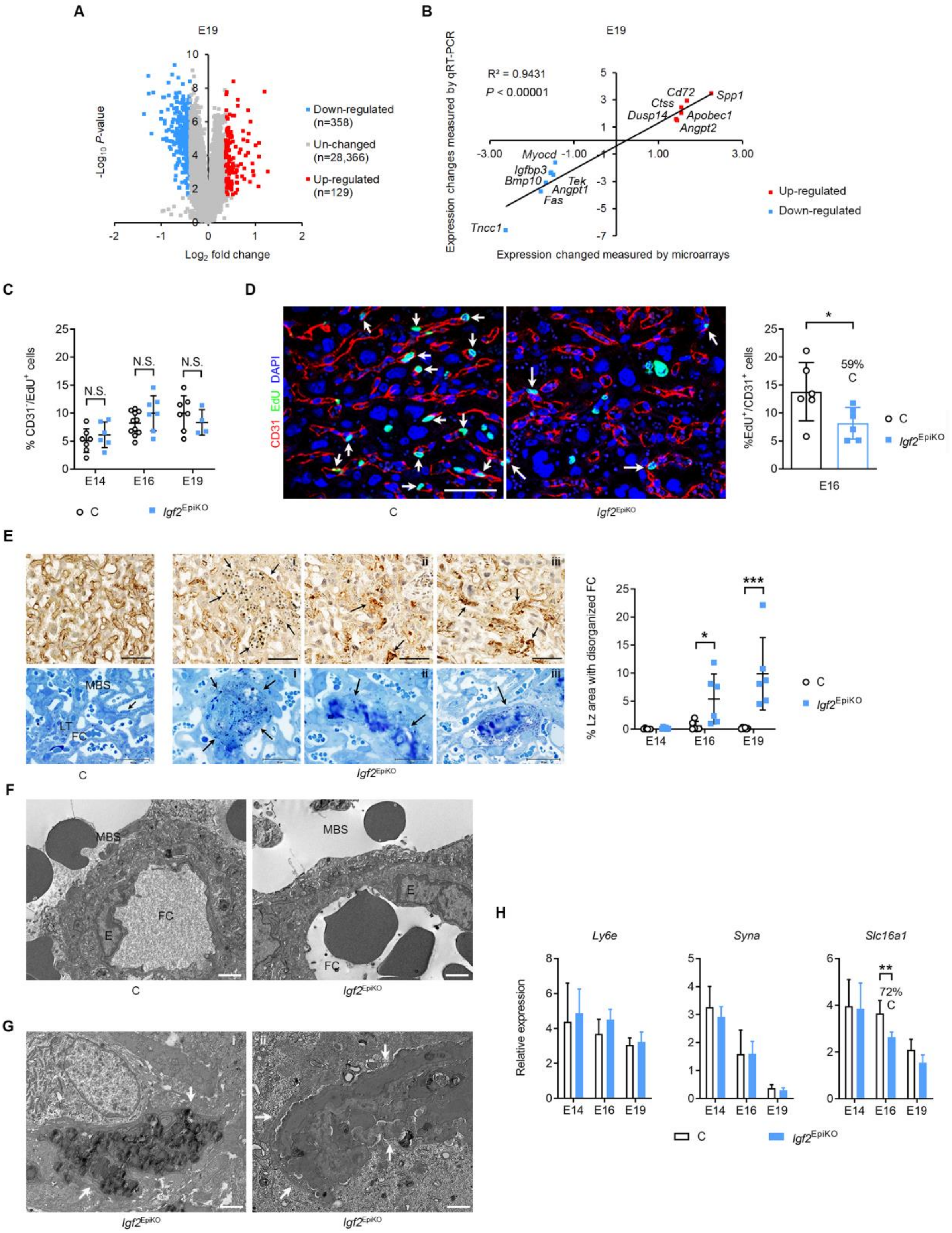


**Figure S4. Fetus-derived *Igf2* deletion (*Igf2*$^{EpiKO}$) alters gene expression in placental Lz and FPEC survival and proliferation.** (**A**) Volcano plot depicting differentially expressed genes (DEG) identified in E19 placental Lz by expression microarray analysis (n=6 samples per genotype, all from male conceptuses). (**B**) Biological validation using qRT-PCR for 13 DEGs (n=6–7 samples per genotype), normalized against three housekeeping genes (*Sdha*, *Gapdh* and *Pmm1*). (**C**) The reduction in proliferation seen in FEPC (Fig. 3e) is not observed in non-endothelial cells from placental Lz measured by flow cytometry analysis after EdU injections (16 hours exposure; n=4–11 per group). (**D**)

Representative confocal microscopy image of FPEC proliferation in control (panel i) versus mutant (panel ii) E16 placentae by immunofluorescent staining for CD31 combined with Click-iT EdU imaging. Arrows point towards FPEC nuclei that incorporated EdU *in vivo* during the 16 hours exposure to the thymidine analogue. Scale bar is 50 µm. The accompanying graph shows data quantification based on counting between 250 to 600 FPEC per sample (n=6 placentae/genotype). **(E)** Top row: CD31 staining in control (C) and *Igf2*$^{EpiKO}$ mutant E16 placental Lz illustrating abnormally large FCs lacking endothelial cells (i) or obstructed capillaries surrounded by fragmented and disorganized FPEC (ii, iii). Scale bars are 50µm. Bottom row: methylene blue-stained E16 placental Lz resin sections (arrows indicate a FPEC in C and thrombotic FC in mutants: i-iii). Scale bars are 30µm. The graph on the right side shows the quantification of percentage of placental Lz areas with disorganized pattern of FC. Measurement of total Lz and abnormal Lz areas was performed using HALO image analysis software (n=6 samples per genotype at each developmental stage). (**F**) Representative transmission electron microscopy (TEM) micrographs showing intact feto-maternal barriers separating fetal capillaries (FC) from maternal blood spaces (MBS) in both controls (C) and *Igf2*$^{EpiKO}$ mutants, at E16. Scale bars are 2µm (n=3 biological replicates per group with 6-15 micrographs/sample). (**G**) TEM micrographs from E16 *Igf2*$^{EpiKO}$ mutants. White arrows point towards an apoptotic endothelial cell (panel i) and a blocked/thrombotic capillary with apoptotic endothelial lining (panel ii). In both cases the overlying trophoblast is intact. Scale bars are 2µm. (**H**) qRT-PCR analysis of genes expressed in Syn-TI in micro-dissected placental Lz in mutants versus controls (n=6–8 samples per group for each developmental time point). For all graphs, data is presented as averages or individual values ± SD; N.S. – non-significant, * $P < 0.05$, ** $P < 0.01$, *** $P < 0.001$ by two-way ANOVA plus Sidak's multiple comparisons tests in (C), (D) and (H) or Mann Whitney test in (E).

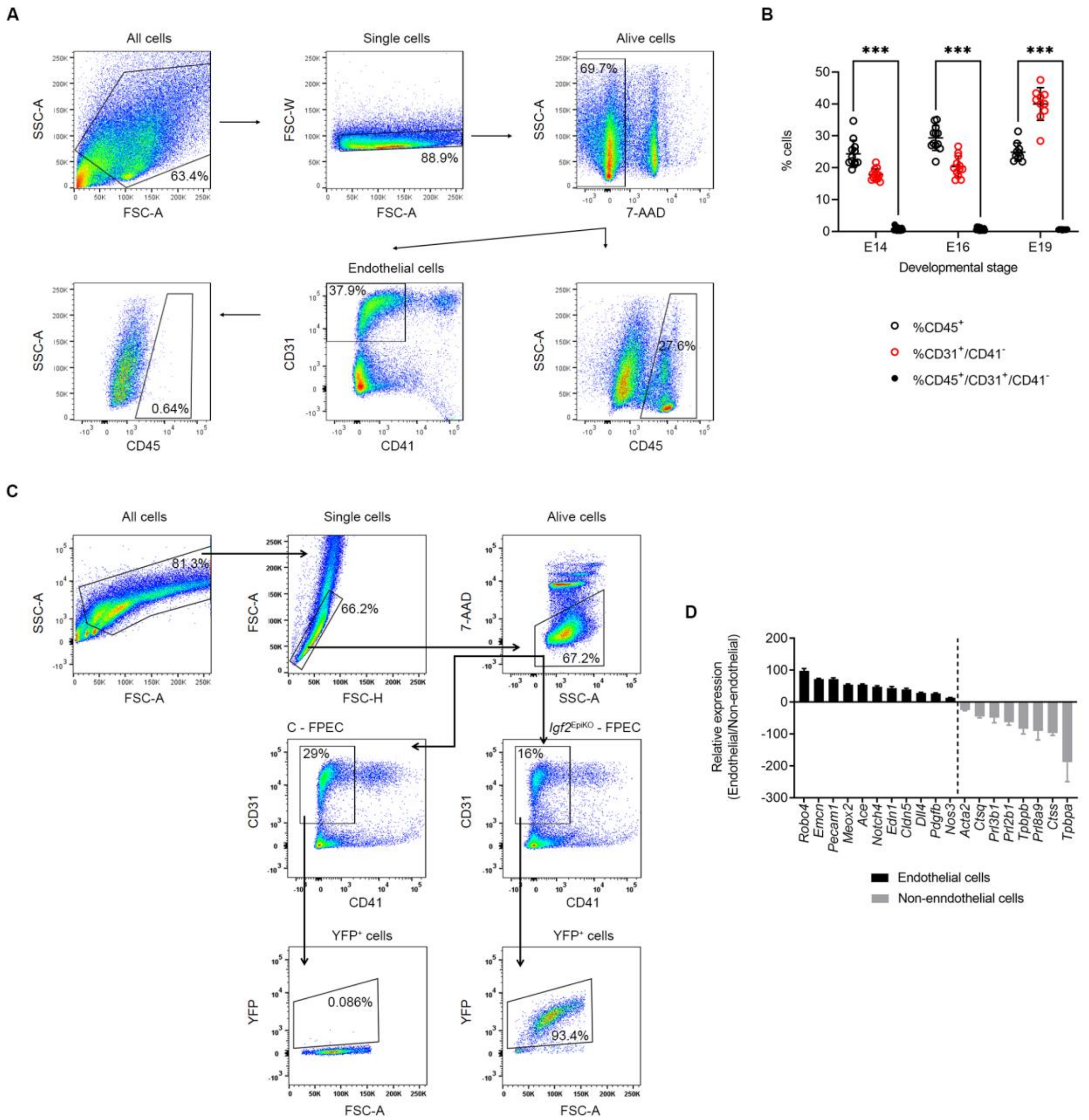


**Figure S5. Experimental design used for FPEC analysis by flow cytometry and FPEC isolation by FACS.** (**A**) Gating strategy used for flow cytometry analysis of CD45 expression (marker of all differentiated hematopoietic cells, except erythrocytes and plasma cells) within FPEC (defined as $CD31^+/CD41^-$ cells). The example shown is from an E19 sample. (**B**) Statistical analysis of CD45+, $CD31^+/CD41^-$ (FPEC) and $CD45^+/CD31^+/CD41^-$ cells at E14, E16 and E19. At all three developmental stages, the proportion of $CD45^+$ within the FPEC ($CD31^+/CD41^-$) is very low. The graph show individual data points with averages and SD (n=10-12 samples from two litters at each developmental stage). *** $p < 0.001$ by two-way ANOVA with Tuckey's multiple comparisons test. (**C**) Gating strategy used for FPEC and YFP analysis by flow cytometry and FPEC isolation by FACS. Mutant FPEC are also positive for YFP (activated by Cre mediated deletion of the *Rosa26* $^{fl}$STOP$^{fl}$YFP STOP cassette), thus providing an internal control for Cre efficiency in each biological sample. (**D**) RNA-seq analysis of marker genes expressed in FPEC or non-endothelial cells isolated by FACS from E16 control placental Lz. The graph shows the relative enrichment in FPEC of known markers of endothelial cells (black) and depletion of marker genes expressed by other cell types found in the placental Lz (grey): pericytes (*Acta2*), sinusoidal trophoblast giant cells (*Ctsq*, *Prl2b1*, *Prl8a9*), parietal trophoblast giant cells (*Prl3b1*) or spongiotrophoblast cells (*Tpbpa*, *Tpbpb*).

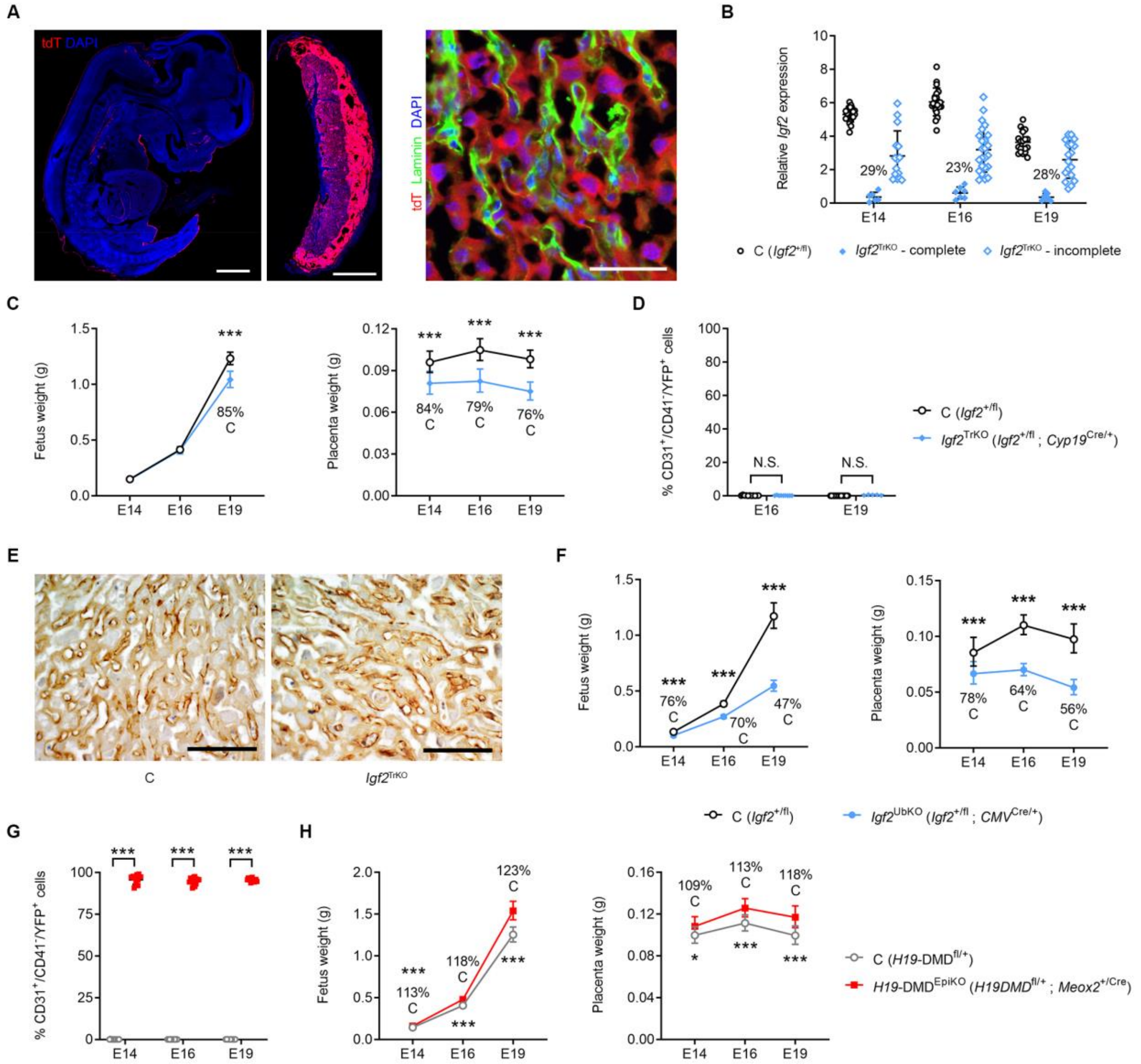


**Figure S6. Specificity and efficiency of *Igf2*$^{TrKO}$, *Igf2*$^{UbKO}$ and *H19*-DMD$^{EpiKO}$ deletions.** (**A**) Representative confocal microscopy on frozen sections from a double transgenic Ai9(RCL-tdT), *Cyp19*$^{Cre}$ fetus and corresponding placenta, at E16, demonstrating high Cre activity (red) in placenta and weak activity in embryonic skin and eye lenses. Right panel: *Cyp19*$^{Cre}$ is only active in the trophoblast cells in placental Lz, as demonstrated by lack of overlapping between the tomato protein (red) and laminin (green) expressed in FPEC. Scale bars are 1 mm (left and middle panel) or 50 µm (right panel). (**B**) Efficiency of *Igf2* deletion by *Cyp19*$^{Cre}$ in *Igf2*$^{TrKO}$ mutants (*Igf2*$^{+/fl}$; *Cyp19*$^{Cre/+}$) versus controls (*Igf2*$^{+/fl}$) evaluated using qRT-PCR in micro-dissected placental Jz layer (n=20–31 samples per genotype). Only 23-29% of all *Igf2*$^{TrKO}$ mutants have high levels of deletion (>80%). (**C**) Placenta growth restriction precedes fetal growth restriction in *Igf2*$^{TrKO}$ mutants (n=4–9 litters at each developmental stage; only mutants with >80% deletion were included in this analysis). (**D**) Flow cytometry analysis showing that *Cyp19*$^{Cre}$ is not expressed in FPEC (note lack of YFP expression in *Igf2*$^{TrKO}$ mutants) (n=6–21 per genotype). (**E**) Representative CD31 immunostainings in E16 control and *Igf2*$^{TrKO}$ placental Lz (scale bars are 100µm) showing no impact of the deletion on FPEC numbers. (**F**) Severe fetal and placental growth restriction in *Igf2*$^{UbKO}$ (*Igf2*$^{+/fl}$; *CMV*$^{Cre/+}$) mutants (n=3-8 litters at each developmental stage). (**G**) Flow cytometry analysis shows that the majority (>80%) of *H19*-DMD$^{EpiKO}$ (*Meox2*$^{+/Cre}$; *H19*-DMD$^{fl/+}$) mutant FPEC express YFP, demonstrating good efficiency of *Meox2*$^{Cre}$ in these cells (n=9–15 per genotype). (**H**) Fetal and placental overgrowth in *H19*-DMD$^{EpiKO}$ mutants (n=3-4 litters at each developmental stage). For all graphs, data is shown as averages or individual values ± SD in (B), (D) and (G) or 95% confidence intervals (95%CI) in (C), (F) and (H). N.S. – non-significant, * $P < 0.05$; ** $P < 0.01$; *** $P < 0.001$ calculated by two-way ANOVA plus Sidak's multiple comparisons tests in (B), (D), and (G) and mixed effects model in (C), (F) and (H).

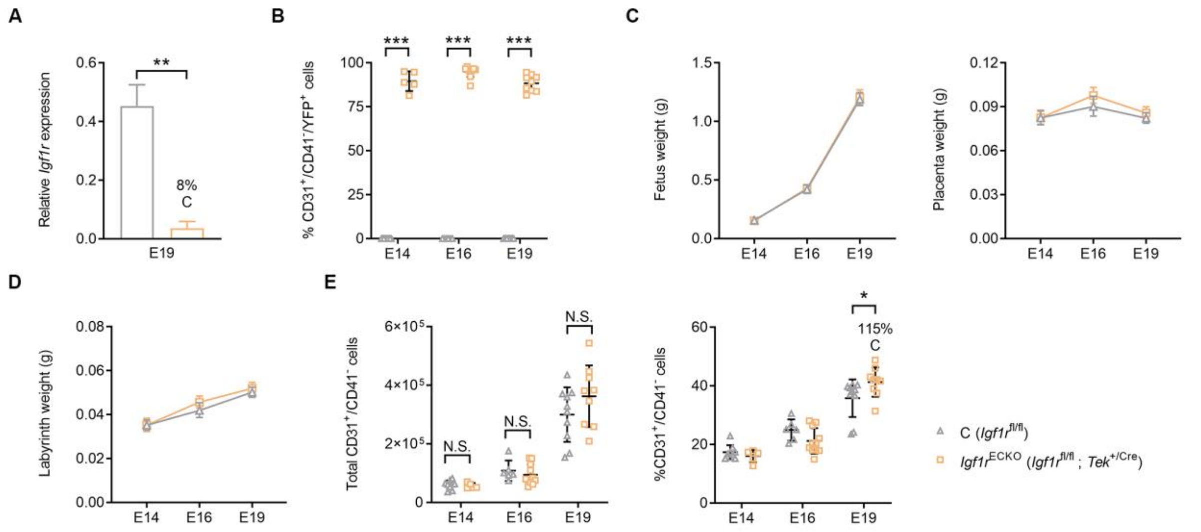


**Figure S7. Conditional deletion of *Igf1r* from endothelium using $Tek^{Cre}$.** (**A**) qRT-PCR analysis of *Igf1r* mRNA levels in primary FPEC isolated by FACS from E19 placental Lz of $Igf1r^{ECKO}$ ($Igf1r^{fl/fl}$; $Tek^{+/Cre}$) mutants versus $Igf1r^{fl/fl}$ controls. (**B**) Flow cytometry analysis showing that the majority (>80%) of $Igf1r^{ECKO}$ mutant FPEC express YFP, demonstrating good efficiency of $Tek2^{Cre}$ in these samples (n=5–11 per genotype). Fetal, placental (**C**) and placental Lz (**D**) growth kinetics are not altered in $Igf1r^{ECKO}$ mutants compared to controls (n=6–18 conceptuses from n=3-7 litters for each developmental stage). (**E**) Total numbers and proportions of FPEC/placental Lz measured by flow cytometry (n=5–11 per genotype). For all graphs, data is shown as individual values or averages ± SD in (A), (B) and (E) or 95%CI in (C) and (D); N.S. – non-significant, * $P < 0.05$, ** $P < 0.01$, *** $P < 0.001$ by Mann-Whitney tests in (A) or two-way ANOVA plus Sidak's multiple comparisons tests in (B) and (E) or mixed effects model in (C) and (D).

**Table S1**. Top 100 highest expressed genes by RNA-Seq from FACS purified E16 feto-placental endothelial cells. FPKM values represent the average of n=4 independent biological replicates. Paternally expressed and maternally expressed imprinted gene symbols are coloured in blue and red, respectively.

**Table S2**. First sheet: list of differentially expressed genes (DEGs) identified through expression microarray analysis in placental Lz samples micro-dissected from E19 *Igf2*$^{EpiKO}$ mutants (average of n=6) and controls (average of n=6). Upregulated and dowregulated gene symbols are coloured in blue and red, respectively. Second sheet: list of pathways enriched in DEGs, as identified by IPA analysis.

**Table S3**. First sheet: list of differentially expressed genes by RNA-Seq from FACS purified feto-placental endothelial cells of E16 *Igf2*$^{EpiKO}$ mutants (average of n=4) and controls (average of n=4). Upregulated and downregulated gene symbols are coloured in blue and red, respectively. Second sheet: list of differentially expressed genes by RNA-Seq from FACS purified feto-placental endothelial cells of E16 *Igf2*$^{ECKO}$ mutants (average of n=4) and controls (average of n=4). Upregulated and downregulated gene symbols are coloured in blue and red, respectively.

**Table S4. Angiostatic and pro-angiogenic factors produced by feto-placental endothelial cells under the control of fetus-derived IGF2**

| Protein | Expression change | Function (cellular compartment) | Role in angiogenesis | PMID |
|---|---|---|---|---|
| CXCL10 | Up-regulated | Cytokine (extracellular space) | angiostatic | 7537965, 7540647, 8611715, 9064358, 10914483 |
| IL15 | Up-regulated | Cytokine (extracellular space) | angiostatic | 28379958 |
| THBS1 | Up-regulated | | angiostatic | 22553494 |
| ADAMTS1 | Up-regulated | Peptidase (extracellular space) | angiostatic | 12716911, 12814950, 17082774, 22776012 |
| APCDD1 | Up-regulated | Membrane-bound glycoprotein (cellular membrane) | angiostatic | 29154126 |
| IFIT2 | Up-regulated | Interferon-induced protein (cytoplasm) | angiostatic | 26515391 |
| IFI16 | Up-regulated | Transcription regulator (nucleus) | angiostatic | 14729471, 21488755 |
| CCL2 | Up-regulated | Cytokine (extracellular space) | pro-angiogenic/ angiostatic if prolonged expression | 15516694, 16888027, 23329645 |
| EGR1 | Up-regulated | Transcription regulator (nucleus) | pro-angiogenic/ angiostatic if prolonged expression | 10339488, 12872165, 16818645, 27041221 |
| KLF4 | Up-regulated | Transcription regulator (nucleus) | pro-angiogenic/ angiostatic if prolonged expression | 24599951, 27431648, 26823670 |
| GDF15 | Up-regulated | Growth factor (extracellular space) | pro-angiogenic | 21773947, 28831101 |
| HBEGF | Up-regulated | Growth factor (extracellular space) | pro-angiogenic | 15289334, 18925469 |
| SERPINE1 | Up-regulated | Protease inhibitor (extracellular space) | pro-angiogenic | 26180080 |
| PLAT | Up-regulated | Peptidase (extracellular space) | pro-angiogenic | 24601228 |
| ISG20 | Up-regulated | Exonuclease (nucleus) | pro-angiogenic | 29195126 |
| HEY2 | Down-regulated | Transcription regulator (nucleus) | pro-angiogenic | 15107403, 16219802, 22421041 |

**Table S5: Mouse strains and crosses**

| **Paternal genotype** | **Maternal genotype** | **Offspring/embryo genotypes** | **Related to Figure** |
|---|---|---|---|
| *Igf2*$^{+/+}$ | *Igf2*$^{+/+}$ | *Igf2*$^{+/+}$ | Figure 1<br>Figure 2I<br>Figure 6<br>Figures S5A, S5B, S5D |
| *Igf2*$^{fl/fl}$;<br>*Rosa26YFP*-stop$^{fl/fl}$ | *Meox2*$^{+/Cre}$ | C (*Igf2*$^{+/fl}$;<br>*Rosa26YFP*-stop$^{fl/fl}$;<br>*Meox2*$^{+/+}$) | Figures 2B, 2C, 2D, 2J<br>Figure 3<br>Figure 4A<br>Figures 5A, 5C<br>Figures S1C, S1D, S1E, S1F, S1G, S1H<br>Figure S2A<br>Figure S4<br>Figure S5C |
| | | *Igf2*$^{EpiKO}$ (*Igf2*$^{+/fl}$;<br>*Rosa26YFP*-stop$^{+/fl}$;<br>Meox2$^{Cre/+}$) | |
| *Igf2*$^{fl/fl}$;<br>*Rosa26YFP*-stop$^{fl/fl}$ | *Tek*$^{+/Cre}$ | C (*Igf2*$^{+/fl}$;<br>*Rosa26YFP*-stop$^{fl/fl}$;<br>*Tek*$^{+/+}$) | Figures 2F, 2G, 2H, 2J<br>Figure 4B<br>Figure 5D<br>Figure S2B<br>Figures S3B, S3C, S3D |
| | | *Igf2*$^{ECKO}$ (*Igf2*$^{+/fl}$;<br>*Rosa26YFP*-stop$^{+/fl}$;<br>Tek$^{Cre/+}$) | |
| *Igf2*$^{fl/fl}$;<br>*Rosa26YFP*-stop$^{fl/fl}$ | *Vav*$^{+/iCre}$ | C (*Igf2*$^{+/fl}$;<br>*Rosa26YFP*-stop$^{fl/fl}$;<br>*Vav*$^{+/+}$) | Figures S3F, S3G, S3H, S3I |
| | | *Igf2*$^{HCKO}$ (*Igf2*$^{+/fl}$;<br>*Rosa26YFP*-stop$^{+/fl}$;<br>*Vav*$^{iCre/+}$) | |
| *Igf2*$^{+/+}$ | *Meox2*$^{+/Cre}$ | *Igf2*$^{+/+}$; *Meox2*$^{Cre/+}$ | Figure S1I |
| | | *Igf2*$^{+/+}$; *Meox2*$^{+/+}$ | |
| *Ai9(RCL-tdT)*$^{fl/fl}$ | *Meox2*$^{+/Cre}$ | *Ai9(RCL-tdT)*$^{+/fl}$ ; *Meox2*$^{Cre/+}$ | Figure S1B |
| | | *Ai9(RCL-tdT)*$^{+/fl}$ ; *Meox2*$^{+/+}$ | |
| *Ai9(RCL-tdT)*$^{fl/fl}$ | *Tek*$^{+/Cre}$ | *Ai9(RCL-tdT)*$^{+/fl}$ ; *Tek*$^{Cre/+}$ | Figures 2E, S3A |
| | | *Ai9(RCL-tdT)*$^{+/fl}$ ; *Tek*$^{+/+}$ | |
| *Igf2*$^{+/+}$ | *Tek*$^{+/Cre}$ | *Igf2*$^{+/+}$; *Tek*$^{Cre/+}$ | Figure S3E |
| | | *Igf2*$^{+/+}$; *Tek*$^{+/+}$ | |
| *Igf2*$^{fl/fl}$;<br>*Rosa26YFP*-stop$^{fl/fl}$ | *Cyp19*$^{+/Cre}$ | C (*Igf2*$^{+/fl}$;<br>*Rosa26YFP*-stop$^{+/fl}$<br>*Cyp19*$^{+/+}$) | Figure 4C<br>Figures S6B, S6C, S6D, S6E |
| | | *Igf2*$^{TrKO}$ (*Igf2*$^{+/fl}$;<br>*Rosa26YFP*-stop$^{+/fl}$<br>*Cyp19*$^{Cre/+}$) | |
| *Ai9(RCL-tdT)*$^{fl/fl}$ | *Cyp19*$^{+/Cre}$ | *Ai9(RCL-tdT)*$^{+/fl}$ ; *Cyp19*$^{Cre/+}$ | Figure S6A |
| | | *Ai9(RCL-tdT)*$^{+/fl}$ ; *Cyp19*$^{+/+}$ | |
| *Igf2*$^{fl/fl}$;<br>*Rosa26YFP*-stop$^{fl/fl}$ | *CMV*$^{+/Cre}$ | C (*Igf2*$^{+/fl}$;<br>*Rosa26YFP*-stop$^{+/fl}$<br>*CMV*$^{+/+}$) | Figure 4D<br>Figure S6F |
| | | *Igf2*$^{UbKO}$ (*Igf2*$^{+/fl}$; | |

| | | | |
|---|---|---|---|
| | | *Rosa26YFP*-stop$^{+/fl}$ *CMV*$^{Cre/+}$) | |
| *Meox2*$^{+/Cre}$ | *H19-DMD*$^{fl/fl}$; *Rosa26YFP*-stop$^{fl/fl}$ | C (*H19-DMD*$^{fl/+}$; *Rosa26YFP*-stop$^{fl/+}$; *Meox2*$^{+/+}$) | Figure 4E<br>Figures S6G, S6H |
| | | *H19-DMD*$^{EpiKO}$ (*H19-DMD*$^{fl/+}$; *Rosa26YFP*-stop$^{fl/+}$; *Meox2*$^{+/Cre}$) | |
| *Tek*$^{+/Cre}$ | *Igf2r*$^{fl/fl}$; *Rosa26YFP*-stop$^{fl/fl}$ | C (*Igf2r*$^{fl/+}$; *Rosa26YFP*-stop$^{fl/+}$; *Tek*$^{+/+}$) | Figures 7A, 7B, 7C, 7D, 7E, 7F, 7G |
| | | *Igf2r*$^{ECKO}$ (*Igf2r*$^{fl/+}$; *Rosa26YFP*-stop$^{fl/+}$; *Tek*$^{+/Cre}$) | |
| *Igf1r*$^{+/fl}$; *Rosa26YFP*-stop$^{+/fl}$ ; *Tek*$^{Cre/+}$ | *Igf1r*$^{fl/fl}$; *Rosa26YFP*-stop$^{fl/fl}$ | C (*Igf1r*$^{fl/fl}$; *Rosa26YFP*-stop$^{fl/fl\ or\ +/fl}$; *Tek*$^{+/+}$) | Figure S7 |
| | | *Igf1r*$^{ECKO}$ (*Igf1r*$^{fl/fl}$; *Rosa26YFP*-stop$^{fl/fl\ or\ +/fl}$; *Tek*$^{+/Cre}$) | |
| | | *Het deletions* (*Igf1r*$^{+/fl}$; *Rosa26YFP*-stop$^{fl/fl\ or\ +/fl}$; *Tek*$^{+/Cre}$) – not used | |

**Table S6. Primers**

| Primers used for genotyping by PCR | | | | | |
|---|---|---|---|---|---|
| **Strain** | **Primer** | **Sequence** | **Primer** | **Sequence** | **Amplicon (bp)** |
| *Igf2*$^{fl/fl}$ | F | TTACAGTTCAAAGCCACCACG | RW<br>RD | GCCAAAGAGATGAGAAGCACC<br>GCCAAACACAGTAAAAAGAAATGC | WT: 324<br>fl: 449<br>del: 384 |
| *Rosa26* $^{fl}$*STOP*$^{fl}$*YFP* | F | TGTTATCAGTAAGGGAGCT | R-WT<br>R-fl | CACACCAGGTTAGCCTTTA<br>AAGACCGCGAAGAGTTTGT | WT: 239<br>fl: 301 |
| *Meox2*-Cre | F | GGACCACCTTCTTTTGGCTTC | R-WT<br>R-Cre | AAGATGTGGAGAGTACGGGGTAG<br>CAGATCCTCCTCAGAAATCAGC | WT: 410<br>Cre: 311 |
| *Tek*-Cre | F | TGTAAACAAGAGCGAGTGGA | R-WT<br>R-Cre | AGAGAATGGCGAGAAGTCAC<br>TGAGTGAACGAACCTGGTCG | WT: 240<br>Cre: 610 |
| *Vav-iCre* | F-WT<br>F-Vav | ATGTCTCCAATCCTTGAACACTG<br>GACTACCTCCTGTACCTGCAAG | R-WT<br>R-Vav | GCAGTGGGAGAAATCAGAACC<br>ACTCTGATTCTGGCAATTTCGG | WT: 254<br>Cre: 329 |
| *Cyp19*-Cre | F | GACCTTGCTGAGATTAGATC | R | AGAGAGAAGCATGTTTAGCTGG | Cre: 545 |
| *CMV*-Cre | F | CGAGTGATGAGGTTCGCAAG | R | TGAGTGAACGAACCTGGTCG | Cre: 390 |
| *H19*-DMD$^{fl/fl}$ | F | CAGGCCTGTCCTCACCTGAAC | R | GCCAGCTTGCCTTGGCAACCCCTT | WT: 387<br>fl: 520 |
| *Igf2r*$^{fl/fl}$ | F | CCTTCCCTCCAGGCCGTTAC | R | GGTGAGGTCTCCATCTGAGTACC | WT: 225<br>fl: 259 |
| *Igf1r*$^{fl/fl}$ | F | CTTCCCAGCTTGCTACTCTAGG | R | CAGGCTTGCAATGAGACATGGG | WT: 124<br>fl: 220 |
| **Primers used for qRT-PCR** | | | | | |
| **Gene** | **Primer** | **Sequence** | **Primer** | **Sequence** | **Amplicon (bp)** |
| *Igf2* | F | AGTCCGAGAGGGACGTGTCTA | R | CGGACTGTCTCCAGGTGTCAT | 102 |
| *Angpt1* | F | GAAGCAACTTCTCAACAGACA | R | TTCTTTGTGTTTTCCCTCCATT | 100 |
| *Angpt2* | F | CTTCTACCTCGCTGGTGAAGAG | R | GCTAAAATCACTTCCTGGTTGG | 106 |
| *Tek* | F | GGAGTGGAGTGAAGAACTAGG | R | GTGGAGTCAGTGATGTTGGAGA | 93 |
| *Fas* | F | CTGCAGACATGCTGTGGATCT | R | GCCTCCTCAGCTTTAAACTCTC | 114 |
| *Ctss* | F | AGAGACCCTACCCTGGACTACC | R | GATTCTTTTCCCAGATGAGACG | 109 |
| *Spp1* | F | ACCATGAGATTGGCAGTGATTT | R | GAGCTGCCAGAATCAGTCACTT | 83 |
| *Tnnc1* | F | GATCTCTTCCGCATGTTTGAC | R | TCAATGTCATCTTCCGTAATGG | 107 |
| *Myocd* | F | ATTCCTGTGCACACTGCTGTAA | R | GAGCTTCTTCACCTTTGGTTTG | 96 |
| *Apobec1* | F | GCACACCTGAGGAAACAAAGTC | R | CAGAGTGGGATCAACAGCTACA | 134 |
| *Cd72* | F | CCAAGGAGAACCTGAAAACTGA | R | GCACCTTTCCTGATATGGAATC | 146 |

| | | | | | |
|---|---|---|---|---|---|
| *Dusp14* | F | CTCCCTGGAAATCCTTAGCAC | R | ACCTCTGGAGCTCATGAAGATG | 133 |
| *Bmp10* | F | CTCTACAACAAATTCGCCACAG | R | GAGCCCATTAAAAGTGACTGGT | 108 |
| *Igfbp3* | F | CAGGCAGCCTAAGCACCTAC | R | GGAACTTGGAATCGGTCACTC | 135 |
| *Adgre1* | F | TAGCTGCTCTTCTGATACCCTC | R | CCAACATTCATCTTGTCCCCTC | 145 |
| *Gcm1* | F | CCGCAAGATTTACCTGAGACC | R | GAATAAGCTTCAGGGGTCCATT | 98 |
| *Syna* | F | AGCCCTCTCTGGACAATATTCA | R | CAAGGTGGGAGAAGATATTTGG | 89 |
| *Synb* | F | CAGCTGACACCCTCATTAAACA | R | ATCCAGAAATGGGAATGAAGTG | 122 |
| *Slc16a1* | F | TCGCAGCTTCTTTCTGTAACAC | R | TCATAGTCAGAGCTGGGTTCAA | 102 |
| *Slc16a3* | F | TGCAGAAGCATTATCCAGATCTAC | R | GTATCGATTGAGCATGATGAGG | 99 |
| *Ly6e* | F | ACATGAGAGTCTTCCTGCCTGT | R | TTCTGATCGGTACATGAGAAGC | 91 |
| *Adamts1* | F | CAAAGGACAGGTGCAAGCTC | R | TTGCACACAGACAGAGGTAGAG | 119 |
| *Cxcl10* | F | CGTCATTTTCTGCCTCATCCTG | R | TGATTTCAAGCTTCCCTATGGC | 134 |
| *Thbs1* | F | ATGTACCCATCCAGAGCATCTT | R | GGTTCCAAAGACAAACCTCACA | 125 |
| *Edn1* | F | GACATCATCTGGGTCAACACTC | R | AAGTCTTTCAAGGAACGCTTGG | 86 |
| *Iigp1* | F | ATGATTTGCCCTCCAGCTTTAC | R | ACTGAATATTCCCTTTTCTCATCCT | 117 |
| *Cdkn1a* | F | GAACATCTCAGGGCCGAAAAC | R | CACTTCAGGGTTTTCTCTTGCA | 96 |
| *Hey2* | F | CTGCCAAGTTAGAAAAGGCTGA | R | CTCATGAAGTCTGTGGCAAGAG | 118 |
| *Igf1r* | F | GTTATCCACGACGATGAGTGC | R | AGTCACCGAATCGATGGTTTTC | 150 |
| *Igf2r* | F | GGAAGACACCAGAACCAGACA | R | TGACACTCATCCTCTGGAAGC | 103 |
| *Insr* | F | GAGAGGATGTGAGACGACGG | R | AGCAGTTCTCCAGCTCATGTAG | 149 |
| *Gapdh* | F | ACAACTCACTCAAGATTGTCAGCA | R | ATGGCATGGACTGTGGTCAT | 121 |
| *Sdha* | F | TTCCGTGTGGGGAGTGTATTG | R | ATTCTGCAGCTCCAGGGTCTC | 135 |
| *Pmm1* | F | ATCCGGGAGAAGTTTGTGGAA | R | GCTGTCTTCATCCAGGCTGTC | 144 |
| *Ppia* | F | AAGGGTTCCTCCTTTCACAGAA | R | GATGCCAGGACCTGTATGCTT | 146 |

**Table S7. Conditions used for immunostainings**

| Staining | Antigen retrieval | Blocking | Primary antibody | Secondary antibody |
|---|---|---|---|---|
| IGF2 | Digestion with 1% pronase (Protease from Streptomyces griseus, Sigma – P6911) in 1xPBS for 10 min at 37°C | 15% Donkey serum (Sigma – D9663) in PBS | Goat anti-human IGF2 (1:50, R&D systems AF-292) overnight at 4°C | AF488 Donkey anti-goat (1:200, Jackson ImmunoResearch – 705-546-147), one hour at room temperature (RT) |
| IGF2R | Boiling for 20 min in Tris-EDTA buffer (10 mM Tris, 1 mM EDTA, 0.05% Tween 20, pH 9.0,) | Animal-free blocking solution (Vector – SP-5030) | Rabbit anti-IGF2R (1:400, Cell Signaling 14364) overnight at 4°C | AF594 Donkey anti-rabbit (1:200, Jackson ImmunoResearch 711-546-152), one hour at RT |
| YFP | Autoclaving for 15 min at 121°C in citric acid buffer (10 mM citric acid, pH 6.0, 0.05% Tween 20) | 5% Donkey serum (Sigma – D9663) in PBS | Goat anti-GFP (1:200, Abcam – ab6673) overnight at 4°C | AF488 Donkey anti-goat (1:200, Jackson ImmunoResearch – 705-546-147), one hour at RT |
| CD31 (immune-histochemistry) | Boiling for 30 min in citric acid buffer (10 mM citric acid, pH 6.0, 0.05% Tween 20) | - 3% H2O2 solution (peroxidase inactivation) 30 min at RT;<br>- 10% Goat serum (Sigma – G9023) and 1% BSA in PBS | Rabbit anti-CD31 (1:50, Abcam – ab28364) overnight at 4°C | Goat anti-Rabbit IgG, biotinylated (1:1000, Abcam – ab6720), one hour at room temperature, then Streptavidin-horse radish peroxidase (1:250 Rockland S000-03), one hour at room temperature, then DAB (Dako – K3468), 3-20 minutes at RT |
| CD31 (immune-fluorescence – assay 1) | Boiling for 30 min in citric acid buffer (10 mM citric acid, pH 6.0, 0.05% Tween 20) | 15% Donkey serum (Sigma – D9663) in PBS | Rabbit anti-CD31 (1:50, Abcam – ab28364) overnight at 4°C | AF594 Donkey anti-rabbit (1:200, Jackson ImmunoResearch 711-546-152), one hour at RT |
| CD31 (immune-fluorescence – assay 2) | Boiling for 20 min in Tris-EDTA buffer (10 mM Tris, 1 mM EDTA, 0.05% Tween 20, pH 9.0,) | Animal-free blocking solution (Vector – SP-5030) | Goat anti-CD31 (1:20, R&D – AF3628) overnight at 4°C | NL557-conjugated Donkey Anti-Goat (1:200, R&D – NL001), one hour at RT |
| F4/80 | Heat-induced antigen retrieval in Target Retrieval Solution (pH=6) – Dako S236984-2 | - Bloxall (peroxidase) Blocking Solution – Vector Labs SP-6000;<br>- Animal-Free Blocker - Vector Labs SP-5030 | Rat anti-Mouse F4/80 (1:20, [CI:A3-1] – Bio-Rad MCA497) 1 hour at RT | - Rabbit anti-Rat IgG (H+L) (1:250, Bethyl A110-322A) 1 hour at room temperature;<br>- Anti-Rabbit HRP (ImmPress – Vector Labs MP-7451) 30 min at RT;<br>- DAB (ImmPact DAB Kit - Vector Labs SK-4105) |
| MCT1 | Proteinase K digestion (Dako – S3020) for 3 minutes at room temperature | 15% Donkey serum (Sigma – D9663) in PBS | Chicken anti-MCT1 (1:200, Merk Millipore – AB1286-I) overnight at 4°C | AF488 Donkey anti-chicken (1:200, Jackson ImmunoResearch – 703-546-155), one hour at RT |

| | | | | |
|---|---|---|---|---|
| MCT4 | Proteinase K digestion (Dako – S3020) for 3 minutes at room temperature | 15% Donkey serum (Sigma – D9663) in PBS | Rabbit anti-MCT4 (1:500, Merck Millipore – AB3314P) overnight at 4°C | AF488 Donkey anti-rabbit (1:200, Jackson ImmunoResearch – 711-546-152), one hour at RT |
| Laminin | Proteinase K digestion (Dako – S3020) for 3 minutes at room temperature | 15% Donkey serum (Sigma – D9663) in PBS | Rabbit anti-laminin (1:500, Dako – Z0097) overnight at 4°C | AF488 Donkey anti-rabbit (1:200, Jackson ImmunoResearch – 711-546-152), one hour at RT |
| Lectin | Digestion with 0.04% pepsin (Sigma – 10108057001) in 0.01M HCl, for 10min at 37°C | - 3% methanol (peroxidase inactivation) for 10min at RT;<br>- 2% bovine serum albumin, 1% skimmed dry milk and 0.1% Tween20, for 15min at RT | biotinylated lectin (1:250 isolectin B4, B-1205, Vector Laboratories) for 90min at 37°C | horseradish peroxide-conjugated streptavidin (1:500 Rockland Immunochemicals S000-03 for 60min at RT) followed by DAB (Sigma D3939) for 10min at RT |
| Cytokeratin | Digestion with 0.04% pepsin (Sigma – 10108057001) in 0.01M HCl, for 10min at 37°C | - 3% methanol (peroxidase inactivation) for 10min at RT;<br>- 2% bovine serum albumin, 1% skimmed dry milk and 0.1% Tween20, for 15min at RT | Rabbit anti-pan cytokeratin (1:75 Novus Biologicals nb600-579) overnight at 4°C | alkaline phosphatase-conjugated goat anti-rabbit (1:500 Abcam ab6722) for 60min at RT followed by NBT/BCIP containing levimasole to block endogenous phosphatase (Thermo Fisher Scientific 34070) for 10min at RT |
| EPCAM (frozen sections) | None | 15% Donkey serum (Sigma – D9663) in PBS | Rat anti-mouse CD326/Epcam Clone G8.8 (1:50, BD Biosciences 552370) overnight at 4°C | AF594 Donkey anti-rat (1:250, Thermo Fisher Scientific A-21209) one hour at room temperature |
